# Effect sizes and test-retest reliability of the fMRI-based Neurologic Pain Signature

**DOI:** 10.1101/2021.05.29.445964

**Authors:** Xiaochun Han, Yoni K. Ashar, Philip Kragel, Bogdan Petre, Victoria Schelkun, Lauren Y. Atlas, Luke J. Chang, Marieke Jepma, Leonie Koban, Elizabeth A. Reynolds Losin, Mathieu Roy, Choong-Wan Woo, Tor D. Wager

**Affiliations:** Dartmouth College, Hanover, NH; Weill Cornell Medical College, New York, NY; Emory University, Atlanta, GA; Columbia University, New York, NY; National Center for Complementary and Integrative Health, National Institutes of Health, Bethesda, MD; National Institute of Mental Health, National Institutes of Health, Bethesda, MD; National Institute on Drug Abuse, National Institutes of Health, Baltimore, MD; University of Amsterdam, Amsterdam; INSEAD Fontainebleau & ICM paris, Paris; Department of Psychology, University of Miami, Miami FL; Department of Psychology, McGill University, Montreal, Quebec; Center for Neuroscience Imaging Research, Institute for Basic Science, Suwon, Gyeonggi-do

**Keywords:** effect size, test-retest reliability, multivariate brain signature, evoked pain, fMRI

## Abstract

Identifying biomarkers that predict mental states with large effect sizes and high test-retest reliability is a growing priority for fMRI research. We examined a well-established multivariate brain measure that tracks pain induced by nociceptive input, the Neurologic Pain Signature (NPS). In N = 295 participants across eight studies, NPS responses showed a very large effect size in predicting within-person single-trial pain reports (d = 1.45) and medium effect size in predicting individual differences in pain reports (d = 0.49). The NPS showed excellent shortterm (within-day) test-retest reliability (ICC = 0.84, with average 69.5 trials/person). Reliability scaled with the number of trials within-person, with ≥60 trials required for excellent test-retest reliability. Reliability was tested in two additional studies across 5-day (N = 29, ICC = 0.74, 30 trials/person) and 1-month (N = 40, ICC = 0.46, 5 trials/person) test-retest intervals. The combination of strong within-person correlations and only modest between-person correlations between the NPS and pain reports indicate that the two measures have different sources of between-person variance. The NPS is not a surrogate for individual differences in pain reports but can serve as a reliable measure of pain-related physiology and mechanistic target for interventions.

**Significance statement:** Current efforts towards translating brain biomarkers require identifying brain measures that can strongly and reliably predict outcomes of interest. We systematically examined the performance of a well-established brain activity pattern, the Neurological Pain Signature (NPS), in a large and diverse sample of participants. The NPS showed excellent reliability, and the reliability scaled with the number of trials within-person. The NPS responses showed strong correlations with pain reports at the within-person level but only modest correlations at the between-person level. The findings suggest that the NPS is not a surrogate for individual differences in pain reports but can serve as a reliable measure of a pain-related physiological target.

**Author Note:** This project was supported by grants R01MH076136 (T.D.W.), R01DA046064, R01EB026549, and R01DA035484. Elizabeth A. Reynolds Losin was supported by a Mentored Research Scientist Development award from National Institute On Drug Abuse of the National Institutes of Health (K01DA045735). Lauren Y. Atlas was supported in part by funding from the Intramural Research Program of the National Center for Complementary and Integrative Health. Yoni K. Ashar was supported by NCATS Grant # TL1-TR-002386. The content is solely the responsibility of the authors and does not necessarily represent the official views of the National Institutes of Health. Code for all analyses and figures is available at https://github.com/XiaochunHan/NPS_measurement_properties. Data for all analyses and figures is available at https://osf.io/v9px7/.

---

Effect sizes and test-retest reliability of the fMRI-based Neurologic Pain Signature

Understanding individual differences in brain activity and their links with behavior is a primary focus of fMRI research. One approach is to establish structure-function associations and make inferences about the brain bases of individual differences. A distinct but related approach uses brain measures to develop biomarkers that can contribute to measuring external constructs (e.g., pain, the risk for mental illness) and inform diagnosis and treatment (FDA-NIH Biomarker Working Group, 2016). For example, research on neural mechanisms of physiological pain helps to understand which brain areas are involved in constructing different kinds of pain experiences in different populations. Meanwhile, the research helps to develop biomarkers that can subtype pain based on pathophysiology, predicting risk for future pain, and more, leading to new ways of understanding, diagnosing, and treating pain. Both establishing structure-function associations and developing biomarkers require brain measures with good measurement properties, including large effect sizes in predicting external variables (e.g., behavior) and high reliability.

Historically, the measurement properties of fMRI activities have not been carefully or frequently assessed. Effect sizes in predicting external variables are calculated at both within-person and between-person levels when repeated measures are collected within each person. Predictions at these levels can be inconsistent due to different sources of variance (Bakdash & Marusich, 2017; Kievit et al., 2013). For example, two variables can be positively correlated at the within-person level while having no relation at the between-person level.

Assessing effect sizes at both within-person and between-person levels prevents incorrect interpretations of the predictions and facilitates a deeper understanding of the brain measures. Test-retest reliability is one type of reliability index, usually measured with an intraclass correlation coefficient (ICC, Shrout & Fleiss, 1979), that assesses temporal stability under repeated tests. Both effect sizes and test-retest reliability rely on a low random error in the measurement. Test-retest reliability also relies on high inter-individual variability, indicating differentiable measures across subjects (Barnhart et al., 2007).

As translational goals accelerate and sample sizes increase, measurement properties of fMRI studies are increasingly a focus of attention (Bennett & Miller, 2010; Button et al., 2013; Dubois & Adolphs, 2016; Elliott et al., 2019, 2020; Hedge et al., 2018; Herting et al., 2018; Kraemer, 2014; Nichols et al., 2017; Noble et al., 2019; O’Connor et al., 2017; Poldrack et al., 2017; Xu et al., 2016; Zuo et al., 2019; Zuo & Xing, 2014). Studies of traditional univariate brain measures provide a pessimistic picture of task fMRI’s measurement properties. Effect sizes of univariate brain measures in local brain regions have often been limited to moderate effect sizes (i.e., Cohen’s d values centered on approximately d = 0.5; Poldrack et al., 2017). The reliability of univariate brain measures in many studies with small samples varies substantially (Letzen et al., 2016; Manuck et al., 2007; Nord et al., 2017; Plichta et al., 2012). A recent meta-analysis of fMRI literature across diverse tasks generally demonstrated low reliability (ICCs < 0.4) of the average activation level of single brain regions of interest (ROI), which did not decrease with longer test-retest interval (Elliott et al., 2020). Similarly, univariate-style approaches to resting-state fMRI studies have found low test-retest reliability, with ICCs < 0.3 at the individual edge-level connectivity (Noble et al., 2019; Pannunzi et al., 2017).

An important trend in the fMRI studies is the development of *a priori* multivariate brain measures that can be used as biomarkers, also called ‘neuromarkers’ or ‘signatures’ (Abraham et al., 2017; Arbabshirani et al., 2017; Doyle et al., 2015; Gabrieli et al., 2015; Haynes, 2015; Kragel et al., 2018; Orrù et al., 2012; Woo, Chang, et al., 2017). Such models consist of patterns of brain activity, connectivity, and other derived features (e.g., graph-theoretic measures) within and across brain regions, which can be applied prospectively to new samples or participants. Because they are pre-specified models applied to new samples without re-fitting, neuromarkers provide an opportunity to evaluate measurement properties across different samples and contexts systematically. Multivariate brain signatures can yield measures with much larger effect sizes (Cohen’s d > 2; Chang et al., 2015; Geuter et al., 2020; Krishnan et al., 2016; Wager et al., 2013; Zunhammer et al., 2018). They also show enhanced test-retest reliability for both task-evoked (ICCs > 0.7; Kragel et al., 2020; Woo & Wager, 2016) and resting-state (ICCs > 0.6; Gordon et al., 2017; Gratton et al., 2020; Yoo et al., 2019; Zuo & Xing, 2014) fMRI measures in some studies. However, this has rarely been assessed across diverse samples and scanners, particularly with respect to a systematic evaluation of effect sizes for within-person and between-person prediction of external variables and test-retest reliability.

In the current study, we evaluated a well-established multivariate brain-based model in the pain domain, i.e., the Neurologic Pain Signature (NPS; Wager et al., 2013). NPS consists of interpretable and stable patterns across brain regions known to show increased activity in pain-related studies. These regions included the thalamus, the posterior and middle insula, the secondary somatosensory cortex, the anterior cingulate cortex, the periaqueductal gray matter, and other regions (see **Figure 1(A)**). The NPS predicts subjective pain intensity in response to noxious thermal (Wager et al., 2013), mechanical (Krishnan et al., 2016), electrical (Krishnan et al., 2016; Ma et al., 2016), and visceral stimuli (Van Oudenhove et al., 2020). In addition, it does not respond to non-noxious warm stimuli (Wager et al., 2013), threat cues (Krishnan et al., 2016; Ma et al., 2016; Wager et al., 2013), social rejection-related stimuli (Wager et al., 2013), vicarious pain (Krishnan et al., 2016), or aversive images (Chang et al., 2015). The NPS provides a neuromarker of a basic mental process with negative affective components, which can serve as an intermediate phenotype potentially relevant to various disorders. For example, enhanced NPS responses, combined with another brain signature related to non-painful sensory processing, discriminated fibromyalgia from pain-free controls with 93% accuracy (López-Solà et al., 2017).

**Figure 1.**
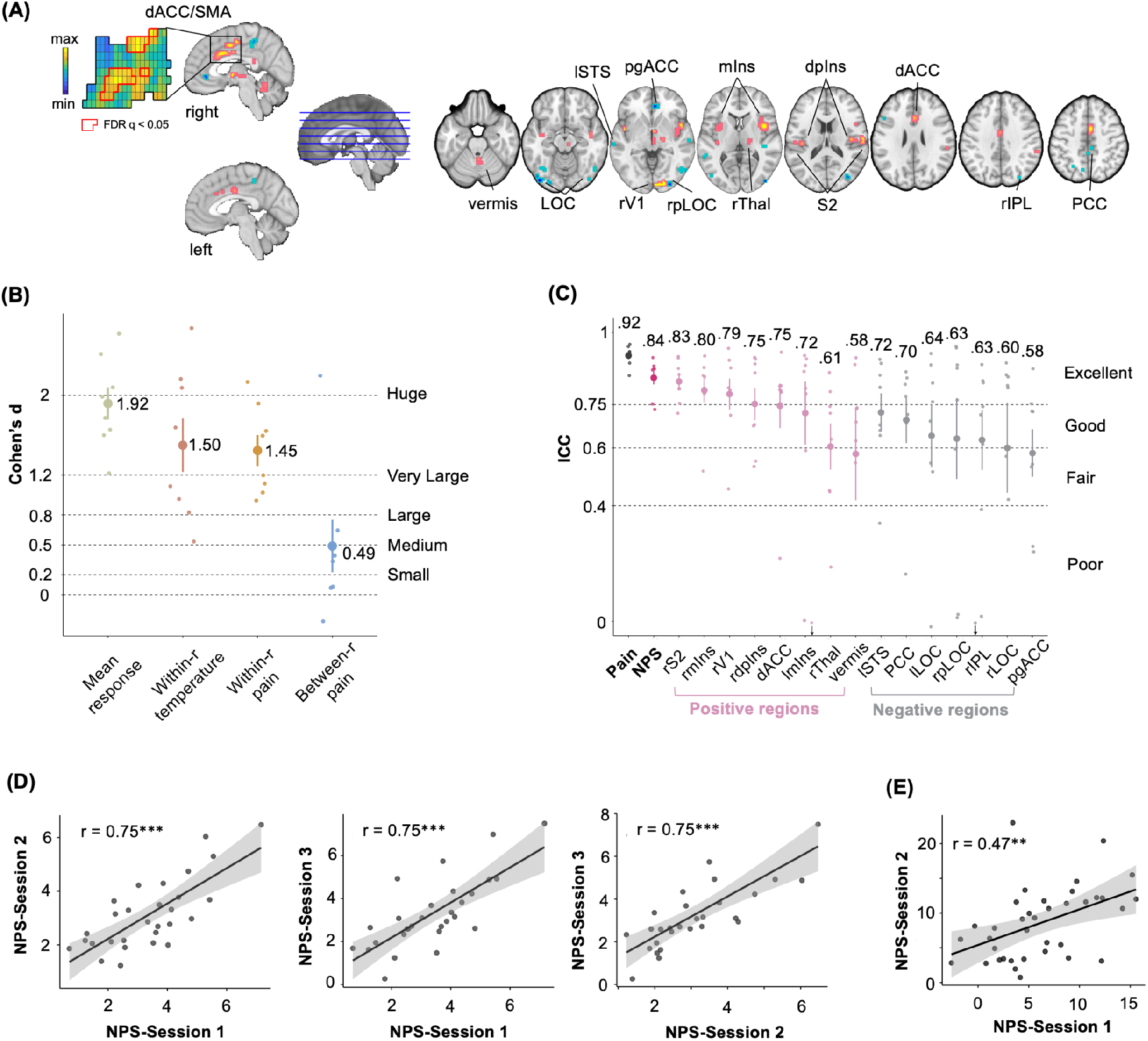
NPS pattern and measurement properties. **(A) The multivariate brain pattern of NPS**. The map shows thresholded voxel weights (at q<0.05 false discovery rate (FDR)) for display only; all weights were used in the subsequent analyses. An example of dACC/SMA unthresholded patterns is presented in the insets; small squares indicate individual voxel weight. Ins denotes Insula, V1 primary visual area, S2 secondary somatosensory cortex, ACC anterior cingulate cortex, Thal thalamus, STS superior temporal sulcus, PCC posterior cingulate cortex, LOC lateral occipital complex, and IPL inferior parietal lobule. Direction is indicated with preceding lowercase letters as follows: r denotes right, l left, m middle, d dorsal, p posterior, pg perigenual. **(B) Four types of NPS effect size**. Each big dot represents a type of averaged effect size of studies 1 to 8; the vertical bar represents the standard error; each small dot represents the effect size of one study. See Figure S1 for the tests of each study. See Figure S2 for the effect sizes of local regions of the NPS. **(C) Short-term test-retest reliability of subjective pain reports, NPS, and local regions.** Each big dot represents the mean reliability of studies 1 to 8; the vertical bar represents the standard error; each small dot represents the reliability of one study. The downward-pointing arrows indicate ICC < 0. See Figure S3 for the illustration of short-term test-retest reliability of the NPS and subjective pain reports. **(D) Illustration of longer-term test-retest reliability of NPS with a 5-day interval**. Correlations of the NPS responses between session 1, session 2 and session 3 in study 9 (ICC = 0.73). Each dot represents one participant; the line represents the linear relationship between the NPS response in sessions 1, 2 and 3, and the shadow represents the standard error. **(E) Illustration of longer-term test-retest reliability of NPS with a 1-month interval**. Correlation of the NPS responses between session 1 and session 2 in the treatment-as-usual control group of study 10 (ICC = 0.46). Each dot represents one participant; the line represents the linear relationship between the NPS response in sessions 1 and 2, and the shadow represents the standard error. *** p < 0.001; ** p<0.005.

NPS effect sizes have been mainly assessed on within-person correlations with pain (Lindquist et al., 2017) and reliability has only been assessed in a preliminary fashion (Kragel et al., 2020; Woo & Wager, 2016). The measurement properties of individual brain regions of the NPS have not been assessed systematically. Comparing the measurement properties of the whole NPS and individual brain regions could help clarify whether NPS’s performance exceeds individual brain regions and reveal the different performances of different individual brain regions. Further, the properties that influence test-retest reliability of the NPS (e.g., amount of data collected per person) have not been systematically examined in detail across studies. Examining these properties could both help understand the NPS as a test case and reveal principles underlying the sources of error and reliability of task fMRI more broadly.

## Methods

We tested four types of effect size for both NPS and local brain regions of interest by analyzing painful stimulus-evoked fMRI and pain reports across ten studies (total N = 444), none of which were used to train the NPS model. The main analyses were conducted on single trial-level data from a multi-study dataset across eight studies (N = 295). In this dataset, we also tested short-term (i.e., within one day) test-retest reliability and several factors potentially influencing it. These factors include the number of trials used, the noxious stimulus intensity, and whether the NPS was applied to pain-versus-rest or a contrast between high and low painful stimulus intensity (Bennett & Miller, 2010, 2013). Studies 9 and 10 evaluated test-retest reliability across 5-days (N = 29) and one-month (N = 120) intervals. We compared the short-term, and longer-term test-retest reliability controlling for the number of trials averaged when calculating the NPS response.

### Datasets description

We analyzed three datasets: (1) a single-trial dataset included 15,940 single-trial images of fMRI activity from healthy subjects with multiple levels of noxious heat and pain ratings within one scan session (i.e., one day) across 295 participants from 8 studies (i.e., study 1 to 8); (2) a study (study 9) with healthy subjects during heat pain tasks with behavioral and fMRI data collected across three sessions with five-days intervals between each session (N = 29); (3) a study (study 10) with chronic back pain subjects receiving pressure pain stimulations with behavioral and fMRI data collected across two sessions with an average of one month between them (N = 120). Participants received a series of painful stimuli and rated their individually experienced pain following each stimulus in all studies. Each study also included psychological manipulation (except for study 3), such as cue-induced expectation and placebo treatment. Descriptive data on age, sex, and other study sample features are given in **Table 1**. The number of trials, stimulation sites, stimulus intensities and durations varied across studies but were comparable; these variables are summarized in **Table 2**. In the studies included, we examined the test-retest reliability of the NPS and pain ratings irrespective of diverse study-specific features and manipulations, which facilitated our conclusion’s generalizability.

**Table 1.** Study demographics, experiment sessions and prior publications

| Study | N | Gender | Ages, M (SD) | # of Sessions | Interval between Sessions (days) | Prior publications |
| --- | --- | --- | --- | --- | --- | --- |
| Study1 | 33 healthy | 22 F | 27.9 (9.0) | 1 | N/A | (Geuter et al., 2020; Lindquist et al., 2017; Woo et al., 2015; Woo, Schmidt, et al., 2017) |
| Study2 | 28 healthy | 10 F | 25.2 (7.4) | 1 | N/A | (Chang et al., 2015; Geuter et al., 2020; Krishnan et al., 2016; Lindquist et al., 2017; Woo, Schmidt, et al., 2017) |
| Study3 | 93 healthy | 49 F | 28.7 (5.7) | 1 | N/A | (Geuter et al., 2020; Losin et al., 2020) |
| Study4 | 17 healthy | 9 F | 25.5 | 1 | N/A | (Atlas et al., 2010; Geuter et al., 2020; Lindquist et al., 2017; Woo, Schmidt, et al., 2017) |
| Study5 | 50 healthy | 27 F | 25.1 (6.9) | 1 | N/A | (Geuter et al., 2020; Lindquist et al., 2017; Roy et al., 2014; Woo, Schmidt, et al., 2017) |
| Study6 | 19 healthy | 10 F | 25.5 (9.5) | 1 | N/A | (Geuter et al., 2020; Jepma et al., 2018) |
| Study7 | 29 healthy | 16 F | 20.4 (3.3) | 1 | N/A | (Lindquist et al., 2017; Woo, Schmidt, et al., 2017) |
| Study8 | 26 healthy | 11 F | 28 (9.3) | 1 | N/A | (Koban et al., 2019; Lindquist et al., 2017; Woo, Schmidt, et al., 2017) |
| Study9 | 29 healthy | 16 F | 29.9 (9.8) | 3 | Ses 1 to 2: 4.93 (4.57);<br>Ses 2 to 3: 4.79 (2.81); | unpublished |
| Study10 | 120 chronic back pain | 61 F | 42.6 (15.6) | 2 | 25 - 40 | Ashar et al., submitted |

**Table 2.** Stimulation protocol

| Study | Stimulus location | Stimulus Intensity (°C) | Stimulus duration (seconds) | Trials per subject | Other experimental manipulations |
| --- | --- | --- | --- | --- | --- |
| Study1 | Arm | 44.3, 45.3, 46.3, 47.3, 48.3, 49.3 | 12.5 | 97 | Cognitive self-regulation intervention to increase or decrease pain |
| Study2 | Arm, Foot | 46, 47, 48 | 11 | 81 | Combination of painful stimuli with heat-predictive visual cues for low, medium, and high pain |
| Study3 | Arm | 47, 48, 49 | 8 and 11 | 36 | Heat stimuli were intermixed with physically and emotionally aversive sound stimuli |
| Study4 | Arm | 41.1 - 47.1 | 10 | 64 | Combination of painful stimuli with heat-predictive auditory cues |
| Study5 | Arm | 46, 47, 48 | 11 | 48 | Combination of painful stimuli with heat-predictive visual cues and with a placebo manipulation |
| Study6 | Leg | 48, 49 | 1.85 | 70 | Combination of painful stimuli with heat-predictive visual cues |
| Study7 | Arm | 44.7, 46.7 | 10 | 64 | Combination of painful stimuli with intervention for perceived control (making vs. observing cue choice) and expectancy (80% vs. 50% probabilities of low pain) |
| Study8 | Leg | 48, 49, 50 | 1.85 | 96 | Combination of painful stimuli with heat-predictive visual cues and unreinforced social information |
| Study9 | Leg | 46, 47, 48 | 12 | 30 | Combine painful stimuli with neural feedback on suppressing NPS activity |
| Study10 | thumbna il | 4, 7 kg/cm2* | 6 | 5 | Data collected in the context of a randomized controlled trial, including a psychotherapy treatment, placebo treatment, and treatment-as-usual control group |

\*Study 10 delivered pressure rather than thermal stimulation.

Data from the study 1 to 8 have been used in previous publications (see **Table 1**). However, the analyses and findings reported here are novel, and the data used for developing the NPS was not included in the current study to avoid double-dipping (Kriegeskorte et al., 2009). Data from the study 9 and 10 have not been published yet. All participants were recruited from New York City and Boulder/Denver Metro Areas. The institutional review board of Columbia University and the University of Colorado Boulder approved all the studies, and all participants provided written informed consent. Participants’ preliminary eligibility was determined through an online questionnaire, a pain safety screening form, and an MRI safety screening form. Participants with psychiatric, physiological, or pain disorders, neurological conditions, and MRI contraindications were excluded before enrollment. No participants were excluded from the study after screening other than individuals who, upon screening, provided different responses that made them ineligible (e.g., developing a physiological disorder).

### Materials and Procedures

#### Thermal and pressure stimulation

We delivered thermal stimulation to multiple skin sites using a TSA-II Neurosensory Analyzer (Medoc Ltd., Chapel Hill, NC) with a 16 mm Peltier thermode endplate, excepting study 3 using the Pathway ATS model and study 8 with a 32 mm Peltier thermode endplate. Study 10 delivered pressure rather than thermal stimulation, using a custom-built pneumatic device pushing a piston into the left thumbnail. At the end of every trial, participants rated pain intensity on a visual analog scale or a labeled magnitude scale (Bartoshuk et al., 2004).

Thermal stimulation parameters varied across studies, with stimulation temperatures ranging from 44.3 °C to 50 °C and stimulation durations ranging from 1.85 to 12.5 seconds. Most studies applied thermal stimulation to the left volar forearm; study 2 also applied the left foot’s dorsum; study 6 and study 8 applied the stimulation to the lower leg. See **Table 2** for stimulation location, intensity levels, duration, number of trials per subject, and other cognitive manipulations.

### fMRI Analysis

#### Preprocessing

We maintained the preprocessing pipelines from the original published studies despite variations across studies as this will likely reflect the variations in preprocessing steps observed across studies in the literature. In studies 1 to 8, structural T1-weighted images were coregistered to each subject’s mean functional image using the iterative mutual information-based algorithm implemented in SPM (Ashburner & Friston, 2005). They were then normalized to MNI space using SPM. Following SPM normalization, study 4 included an additional step of normalization to the group mean using a genetic algorithm-based normalization (Atlas et al., 2010, 2014; Wager & Nichols, 2003). In each functional run, we removed initial volumes to allow for image intensity stabilization. We also identified image-intensity outliers (i.e., ‘spikes’) by computing the mean and standard deviations (SD, across voxels) of intensity values for each image for all slices to remove intermittent gradient and severe motion-related artifacts present to some degree in all fMRI data. We first computed both the mean and the SD of intensity values across each slice for each image to identify outliers. Mahalanobis distances for the matrix of (concatenated) slice-wise mean and standard deviation values by functional volumes (overtime) were computed. Any values with a significant χ2 value (corrected for multiple comparisons based on the more stringent of either false discovery rate or Bonferroni methods) were considered outliers. In practice, less than 1% of the images were deemed outliers. The outputs of this procedure were later included as nuisance covariates in the first-level models. Next, functional images were corrected for differences in each slice’s acquisition timing and were motion-corrected (realigned) using SPM. The functional images were warped to SPM’s normative atlas (warping parameters estimated from coregistered, high-resolution structural images), interpolated to 2 × 2 × 2 mm^3^ voxels, and smoothed with an 8 mm FWHM Gaussian kernel. This smoothing level has been shown to improve inter-subject functional alignment while retaining sensitivity to mesoscopic activity patterns consistent across individuals (Shmuel et al., 2010).

The preprocessing of study 9 and 10 were conducted using *fMRIPrep* 1.2.4 (Esteban et al., 2019; Esteban, Blair, et al., 2018; RRID:SCR_016216). The BOLD reference was co-registered to the T1w reference. Co-registration was configured with nine degrees of freedom to account for distortions remaining in the BOLD reference. Head-motion parameters with respect to the BOLD reference (transformation matrices, and six corresponding rotation and translation parameters) are estimated. The BOLD time-series were resampled onto their original, native space by applying a single, composite transform to correct for head-motion and susceptibility distortions. The BOLD time-series were resampled to *MNI152NLin2009cAsym* standard space, generating a *preprocessed BOLD* run in *MNI152NLin2009cAsym* space. The preprocessed BOLD runs were smoothed with a 6 mm FWHM Gaussian kernel. We identified image-intensity outliers (i.e., ‘spikes’) using Mahalanobis distances (3 standard deviations) and dummy regressors were included as nuisance covariates in the first level. Besides, twenty-four head motion covariates per run were entered into the first level model as well (displacement in six dimensions, displacement squared, derivatives of displacement, and derivatives squared).

### General linear model (GLM) analyses

For studies 1 to 8, a single trial, or “single-epoch”, design and analysis approach was employed to model the data. Quantification of single-trial response magnitudes was done by constructing a GLM design matrix with separate regressors for each trial, as in the “beta series” approach (Mumford et al., 2012; Rissman et al., 2004). First, boxcar regressors, convolved with the canonical hemodynamic response function (HRF), were constructed to model cue, pain, and rating periods in each study. Then, we included a regressor for each trial, as well as several types of nuisance covariates. Because each trial consisted of relatively few volumes, trial estimates could be strongly affected by acquisition artifacts that occur during that trial (e.g., sudden motion, scanner pulse artifacts). Therefore, trial-by-trial variance inflation factors (VIFs; a measure of design-induced uncertainty due, in this case, to collinearity with nuisance regressors) were calculated, and any trials with VIFs that exceeded 2.5 were excluded from the analyses. Single-trial analysis for study 2 and 4 were based on fitting a set of three basis functions, rather than the standard HRF used in the other studies. This flexible strategy allowed the shape of the modeled hemodynamic response function (HRF) to vary across trials and voxels. This procedure differed from that used in other studies because (a) it maintains consistency with the procedures used in the original publication on study 4 (Atlas et al., 2010), and (b) it provides an opportunity to examine predictive performance using a flexible basis set. For both studies, the pain period basis set consisted of three curves shifted in time and was customized for thermal pain responses based on previous studies (Atlas et al., 2010; Lindquist et al., 2009). To estimate cue-evoked responses for study 4, the pain anticipation period was modeled using a boxcar epoch convolved with a canonical HRF. This epoch was truncated at 8 s to ensure that fitted anticipatory responses were not affected by noxious stimulus-evoked activity. As with the other studies, we included nuisance covariates and excluded trials with VIFs > 2.5. In study 4 we also excluded trials that were global outliers (those that exceeded 3 SDs above the mean). We reconstructed the fitted basis functions from the flexible single-trial approach to compute the area under the curve (AUC) for each trial and in each voxel. We used these trial-by-trial AUC values as estimates of trial-level anticipatory or pain-period activity. For studies 9 and 10, we estimated a GLM for each participant, including the nuisance covariates generated in preprocessing and three regressors of interest: pain stimuli, pain ratings, and button presses, each convolved with the standard HRF.

### Computing Neurologic Pain Signature (NPS) responses

We computed a single scalar value for each trial and each subject, representing the NPS pattern expression in response to the thermal and pressure pain stimulus (using the contrast [Pain Stimulation minus Baseline] images). There are three methods to calculate the NPS pattern response, given the NPS is represented as a vector ****x****, brain response to pain stimulus as a vector ****y,**** and the voxel number in the brain mask as ****n****: (1) dot-product 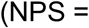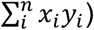, which combine whole-image magnitude and spatial similarity information; (2) cosine similarity 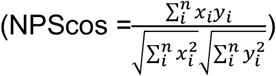, which excludes whole-image magnitude information, representing the dot-product of unit vectors; (3) correlation 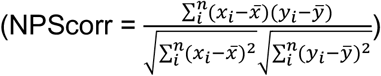 which excludes information related to whole-image mean and magnitude, equivalent to the cosine similarity between centered vectors. The effect size and reliability of these three NPS response metrics were not significantly different from each other (see **Table S5**). Thus we reported the results of the dot product of the NPS in the main text.

To test whether NPS’s performance exceeds individual brain regions within NPS, we also computed the pattern expression, i.e., dot-product, for each brain area within NPS. The individual brain areas were defined based on the NPS map thresholded at q < 0.05 FDR, k > 10. We compared the effect size and the reliability of individual brain regions with the whole NPS pattern using paired t-tests by treating the study as the observation unit and corrected the multiple comparisons using q<0.05 FDR. In most of the regions in the NPS, pain is associated with the increased overall activity, i.e., positive brain regions, including the right middle Insula (rmIns), the right dosal posterior Insula (rdpIns), the left middle Insula (lmIns), the right secondary somatosensory cortex (rS2), the dorsal anterior cingulate cortex (dACC), the right Thalamus (rThal), vermis and the right primary visual area (rV1). Such regions include the major targets of ascending nociceptive afferents. In a subset of other regions, pain is associated with the decreased overall activity, i.e., negative brain regions, including the perigenual ACC (pgACC), the posterior cingulate cortex (PCC), right inferior parietal lobule (rIPL), left lateral occipital complex (lLOC), right posterior lateral occipital complex (rpLOC), right lateral occipital complex (rLOC), and left superior temporal sulcus (lSTS).

These regions are not strongly linked to nociception and are not direct targets of nociceptive afferents; rather, they have been associated with a variety of affective, autonomic, social, self-referential, and decision-making functions (Roy et al., 2012, 2014).

### Effect size analysis

We analyzed four types of effect sizes of the NPS in the single-trial dataset: (1) *Mean response [Pain minus Baseline]*: the mean NPS response across all trials irrespective of the temperature and experiment manipulations. A one-sample t-test was conducted for all participants in each study; (2) *within-person correlation with temperature*: correlation between the temperature and NPS response. A one-sample t-test was conducted for the correlation coefficients of all participants for each study; (3) *Within-person correlation with pain reports*: correlation between pain reports and the NPS response. A one-sample t-test was conducted for the correlation coefficients of all participants for each study. (4) *Between-person correlation with pain reports*. The mean NPS response and mean pain reports of each participant were calculated by the average of each participant’s trials. The correlation between the NPS response and pain reports was calculated across all participants for each study. The effect size was determined by Cohen’s d values, which are commonly characterized as follows: 0.20 indicates small; 0.50 indicates medium; 0.80 indicates large, and 1.20 indicates very large effect size (Cohen, 2013; Sawilowsky, 2009). In between-person correlations, the transformation between r and cohen’s d is 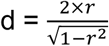.

### Test-retest reliability analysis

Test-retest reliability of the mean NPS response [Pain minus Baseline] was determined by the intra-class correlation coefficient (ICC; Koo & Li, 2016; McGraw & Wong, 1996; Shrout & Fleiss, 1979). To compare with the NPS, we also tested the reliability of the mean pain reports using ICC. ICC is calculated by mean squares obtained through analysis of variance among a given set of measures. We characterized two types of test-retest reliability, i.e., short-term and longer-term test-retest reliability, based on the time interval between measures. In the single-trial dataset, which includes studies 1 to 8, we calculated the short-term test-retest reliability since data were collected within one session. To do so, we constructed a two-way mixed-effects model with time (1st vs. 2nd half trials) as a fixed effect and subjects as a random effect. Since we were interested in the reliability of the averaged measures of the 1st and 2nd half trials (i.e., the average of two halves, k = 2), the mixed-effect model is referred to as ICC(3,k) = (BMS - EMS) / BMS. BMS represents the mean square for between-person measures, and EMS represents the mean square for error. The ICC values in the current study were calculated using the ICC function in the ‘psych’ library in R.

For studies 9 and 10, we assessed the longer-term test-retest reliability since data were collected across sessions with longer time intervals. We also constructed a two-way mixed-effects model with time (multiple sessions) as a fixed effect and subjects as a random effect. Instead of calculating ICC(3,k), we calculated ICC(3,1) = (BMS − EMS) / (BMS + (k − 1) * EMS) for longer-term test-retest reliability since we were interested in the measure of one session, not the average of all sessions. BMS represents the mean square for between-person measures, EMS represents the mean square for error, and k represents the number of scanning sessions (Koo & Li, 2016; McGraw & Wong, 1996; Shrout & Fleiss, 1979).

Measures with ICCs are commonly characterized as follows: less than.40 are thought to have poor reliability, between .40 and .60 fair reliability, .60 and .75 good reliability, and greater than .75 excellent reliability (Cicchetti & Sparrow, 1981). We also reported the 95% confidence interval of ICC values (Koo & Li, 2016; McGraw & Wong, 1996).

## Results

### NPS effect sizes

We tested four types of effect sizes of the NPS in the single-trial dataset. (1) *Mean response [Pain minus Baseline]*: mean responses of the NPS were significantly larger than zero in each of the 8 studies (t = 5.02 − 19.22, ps < 0.001; mean d = 1.92, ranging from 1.22 to 2.62). (2) *Within-person correlation with temperature*: the within-person correlations between the NPS and temperature were significantly larger than zero in each of the 8 studies as well (mean r = 0.05 − 0.42, t = 2.32 − 18.91, ps < 0.05; mean d = 1.50, ranging from 0.53 to 2.67). (3) *Within-person correlation with pain reports*: the within-person correlations between the NPS and subjective pain reports were significantly larger than zero in each of the 8 studies (mean r = 0.14 − 0.35, t = 4.81 − 11.49, ps < 0.001; mean d = 1.45, ranging from 0.94 − 2.13). (4) *Between-person correlation with pain reports*: the between-person correlations between the mean NPS and mean subjective pain rating (i.e., individual differences) were only significant in 1 out of 8 studies (r = −0.13 − 0.74, p = 0.69e-3 − 0.70; mean d = 0.49, ranging from −0.27 to 2.20; see **Figure 1 and Figure S1** for four types of tests and effect sizes; see **Table S1** for the statistical details of each study). In study 4, the only study showed significant between-person correlation, the stimuli were tailored to the individuals to elicit matched subjective pain, whereas the other studies applied the same stimuli for everyone. Thus the individual differences in subjective pain are not stimulus-driven in these studies except for study 4.

To test whether NPS’s performance exceeds individual brain regions within the NPS, we did the same analyses for each local brain area of the NPS and compared the effect sizes with the NPS. Generally, positive brain regions had higher effect sizes than negative brain regions and the effect sizes of the full NPS were the highest in all four tests (see **Figure S2**). To confirm the difference in the effect sizes between NPS and local brain regions, we conducted paired t-tests treating the study as the unit of the observation and corrected the multiple comparisons using q<0.05 FDR. The NPS has (1) significantly larger effect size than most local brain regions in the mean response, except for the rmIns (mean±se = 1.92±0.16 vs. 1.72±0.19); (2) significantly larger effect size in the within-person correlation with the temperature, except for the rmIns (1.50±0.27 vs. 1.21±0.26); (3) significantly larger effect size in the within-person correlation with the subjective pain reports, except for the dACC (1.45±0.16 vs. 1.19±0.12); (4) does not significantly differ in effect size in the between-person correlation with the subjective pain reports from most brain regions, except for the rIPL (0.49±0.26 vs. −0.27±0.17) (see **Table S2** for all statistic details).

### Test-retest reliability

The short-term test-retest reliability of the NPS calculated in the single-trial dataset was distributed from good to excellent among 8 studies (ICC = 0.73 − 0.91; mean±s.e. = 0.84±0.02; see **Table S3** for more details), which was significantly smaller than the reliability of subjective pain reports (ICC = 0.85 − 0.96; mean±s.e. = 0.92±0.01; paired-t test: t(7) = 4.11, p = 0.005). Reliability of the NPS was numerically higher than any local brain regions and was significantly higher than rThal and pgACC (q < 0.05 FDR; see **Figure 1(C)** and **Table S4** for statistical details).

### Most pain-predictive and reliable NPS regions

Among the brain regions in the NPS pattern, we identified several regions with the greatest promise for predicting stable individual differences in pain. Those regions ought to have the highest combination of within-person and between-person correlation with pain reports and test-retest reliability. This is because within-person correlation with pain reports is meaningful in terms of the relationship between NPS and pain reports, which might be driven by factors separate from what drives interindividual differences. Between-person correlation with pain reports is of primary interest for stable individual differences, though the effect sizes are moderate in our results. Besides, reliability is an important precondition for predicting stable individual differences in pain. A combination of three cutoffs, i.e., d > 0.2 for both within-person and between-person correlation with pain, and ICC > 0.6 for short-term test-retest reliability, identified six local regions including lmIns, rmIns, rdpIns, dACC, rS2, and rThal (see **Table S2** and **Table S4**).

### Longer-term test-retest reliability

The longer-term test-retest reliability was tested in studies 9 and 10. For study 9, both reliability of the NPS and pain reports were excellent (ICC = 0.74, 95CI = [0.61, 0.84] and 0.87, 95CI = [0.80, 0.92]; see **Figure 1(D)**). The time interval between session 1 and session 2 was 4.93 ± 4.57 days, and the time interval between session 2 and session 3 was 4.79 ± 2.81 days. Study 10 was a clinical trial randomizing chronic back patients to a psychological treatment, a placebo treatment, or a control group (n = 40 per group), with approximately 1 month between the two assessment sessions. In the control group, the reliability of the NPS was fair (ICC = 0.46, 95CI = [0.22, 0.65]; see **Figure 1(E)**) and the reliability of pain reports was poor (ICC = 0.26, 95CI = [−0.15, 0.49]). The reliabilities of the NPS and pain reports in the psychotherapy group and the placebo group were poor (see **Table S3** for details). Reliability in study 10 was likely limited by the low number of trials (5 trials per person) used in this study (see next section).

### How does the number of trials influence reliability?

We tested how the number of trials of the heat stimuli influences the test-retest reliability. The results in **Figure 2(A)** left panel showed that the more trials averaged to calculate the NPS response, the higher the ICC values in each of the 8 studies. On average, 60 or more trials per condition were required to achieve excellent reliability of the NPS. Given the same number of trials being averaged, ICC values in study 9 (30 trials) and study 10 (5 trials) with longer time intervals were comparable with ICC values of studies 1 to 8. The trend was flatter for the test-retest reliability of subjective pain reports, which achieved an excellent level with even one trial.

**Figure 2.**
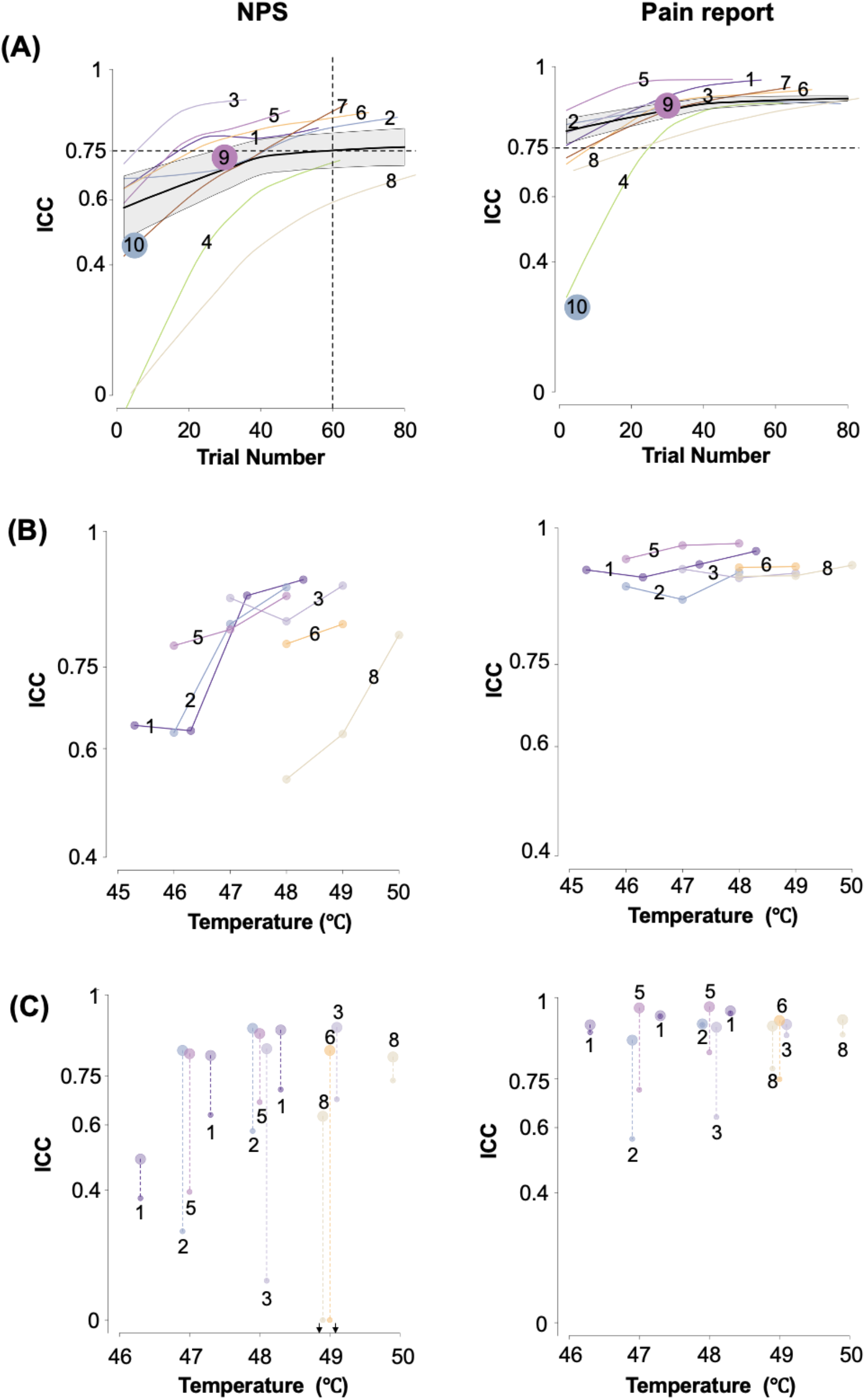
Factors that influence the reliability of the NPS response (left column) and subjective pain reports (right column). The small numbers from 1 to 10 correspond to studies 1 to 10. **(A) The Influence of the trial number and time interval between sessions.** The ICC values were calculated based on different trial numbers. Each line with color shows the nonlinear relationship between the trial number and the ICC values of the corresponding study (fitted using the *loess* function in R). The ICC values estimated with less than 10 participants were excluded due to poor estimation. The black line showed the average of studies 1 to 8, which was weighted by the square root of the number of participants in each study. The grey shadow presents the standard error, which was also weighted by the square root of the number of participants in each study. On average, to achieve excellent reliability, at least 60 trials were required to calculate the NPS response. Reliability was comparable in studies 9 and 10 with a longer time interval across 5-day and 1-month given the same number of trials (trial number = 30 and 5). The reliability of pain reports were excellent in general but were poor in study 10. **(B) The influence of the temperature of the heat stimuli.** Only participants with more than 4 trials in each temperature were included in the ICC calculation. The ICC values estimated with less than 13 participants were excluded due to poor estimation. Under these criteria, the study 4 and 7 were with no ICC value presented in the plot. NPS responses are more reliable in higher temperature stimuli. Whereas pain reports are reliable across all temperature stimuli. **(C) The influence of the types of contrast.** The larger dots represent the ICC values of the measurements calculated by comparing a temperature condition with the baseline, and the smaller dots represent the ICC values of the measurements calculated by comparing a temperature condition with the lowest temperature condition in each study. The length of the dashed line represents the difference between the ICC values of measurements calculated with different types of contrast. The downward-pointing arrow indicates ICC < 0. The measurements calculated by comparing with a control condition are less reliable than by comparing with the implicit baseline in virtually every case.

### How does the effect size of stimuli influence reliability?

The property of the stimulus itself might influence the reliability, such as the effect size it induced. For example, heat stimuli with higher temperatures might generally induce higher pain effects. The results in **Figure 2(B)** left panel showed that NPS responses induced by higher temperature had higher test-retest reliability. However, this was not the case for the subjective pain rating, which was very reliable across all temperatures. NPS responses might be more specific for high painful stimulus intensity, while subjective pain rating could represent a wider range of pain levels in a reliable way.

### How does the type of contrast influence reliability?

There are two commonly used methods to calculate the brain response to an experimental condition, comparing a condition with the implicit baseline or to a control condition. The results in **Figure 2(C)** left panel showed that the reliability of NPS dropped when the response of NPS was calculated in contrast with a lower temperature, instead of the implicit baseline (ICC mean±s.e. = 0.25±0.17 vs. 0.81±0.03, which was calculated by averaging the reliability of all temperatures in one study first and calculating the mean and standard error of the reliability across all studies. Same below.). The drop of the reliability was smaller in subjective pain reports (ICC mean±s.e. = 0.80±0.03 vs. 0.93±0.01). This finding indicates that using a contrast with a control condition with low reliability could reduce the reliability of the contrast measure.

## Discussion

Current efforts towards the translation of brain biomarkers have renewed interest in brain measures’ effect sizes in predicting outcomes of interest and reliability. With a large effect size, a measure can be diagnostic of outcomes at the individual level (Poldrack et al., 2017; Reddan et al., 2017). Test-retest reliability is a prerequisite for stable prediction of individual differences (Bennett & Miller, 2010; Drost & Others, 2011; Nakagawa & Schielzeth, 2010; Streiner, 2003). We systematically evaluated the effect sizes and test-retest reliability of the NPS across ten studies and 442 participants. The NPS showed a very large effect size in predicting within-person single-trial pain reports (mean d = 1.45, ranging from 0.94 to 2.13). The effect size in predicting individual differences in pain reports is medium and heterogeneous across studies (mean d = 0.49, ranging from −0.27 to 2.20). The NPS showed excellent short-term (within-day) test-retest reliability (mean ICC = 0.84). Reliability was comparable in a study with a longer time interval across 5-day (N = 29, ICC = 0.74). It was lower in a study with 1-month test-retest intervals (N = 40, ICC = 0.46), though this may have been driven by the low number of trials (5 trials per person) rather than the longer time interval.

The current findings with a large sample of participants indicate that the NPS measures neurophysiological processes related to evoked pain with large effect sizes at the within-person level and high test-retest reliability. However, as a measure of individual differences in pain sensitivity, the NPS does not reduplicate pain reports. The NPS is only modestly related to the pain reports. This inconsistency of the effect sizes at within-person and between-person levels could be led by the different sources of variance underlying the NPS responses and pain reports (see **Figure 3(A)**). At the within-person level, different temperatures across trials are among the primary sources of variance in NPS responses and pain reports. The effect sizes of within-person correlations between the NPS and the temperatures were distributed from medium to huge (d = 0.53 − 2.67). The effect sizes of within-person correlations between the pain reports and the temperatures were distributed from very large to huge (d = 1.58 − 12.41). Both the NPS and pain reports are responsive to noxious stimuli intensities.

**Figure 3.**
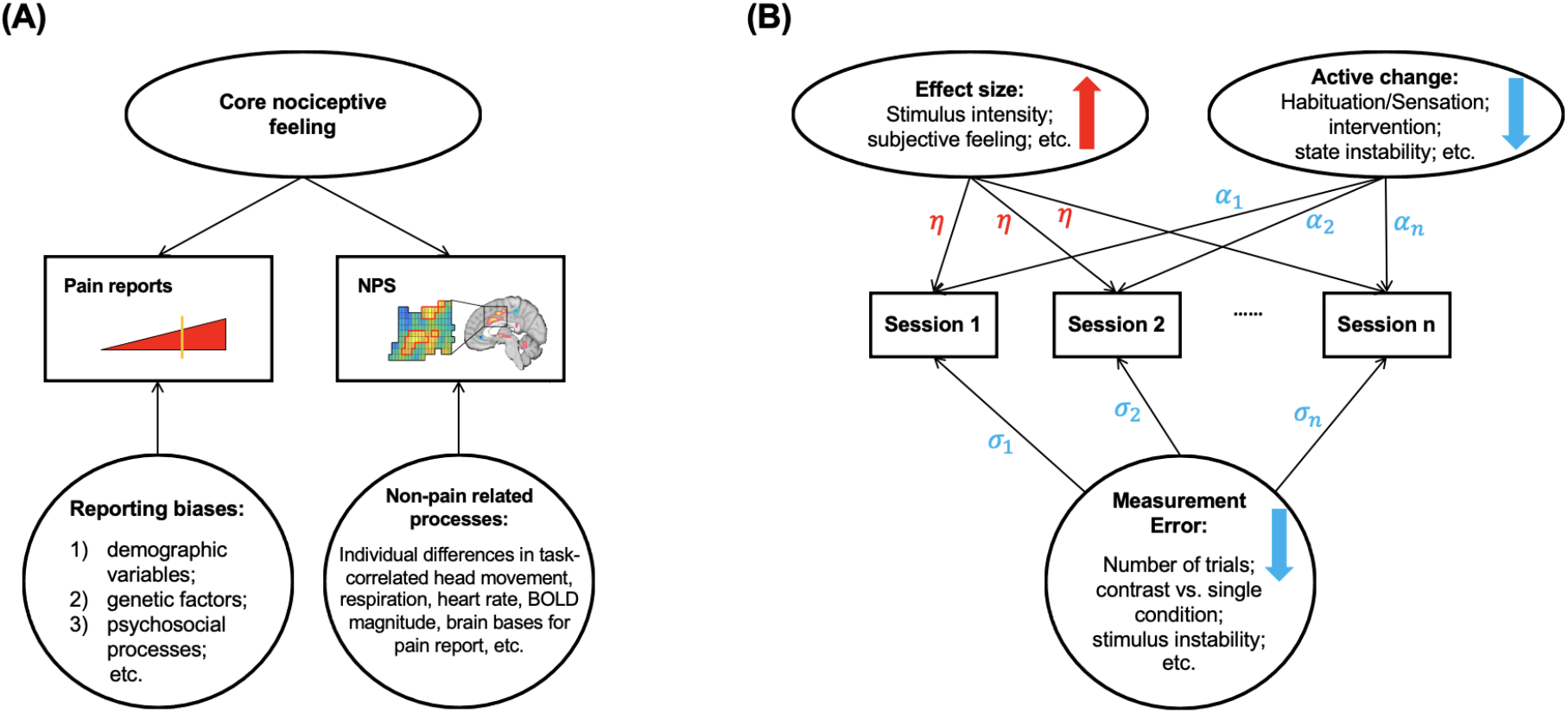
Summary of variances and factors that influence the effect size and reliability. **(A) Different sources of variance at the between-person level for the NPS and self-report pain.** Rectangles represent the observed variables, i.e., pain reports and NPS. Ellipses represent the latent variables that we aim to measure, i.e., the core nociceptive feeling. The circle represents sources of variance that add to each observed measure. Both pain reports and NPS activity measure the core nociceptive circuits that generate pain experience. However, different sources of variance at the between-person level reduce the correlation between pain reports and the NPS response. It suggests that the NPS is not as useful as a surrogate measure for pain reports. In contrast, the NPS could be useful as an objective biological target to measure physiological pain in combination with subjective pain reports. **(B) Factors that influence reliability.** Rectangles represent the observed variables, such as the NPS response, across different sessions. Ellipses represent the latent variables that we are interested in modeling. Results suggest that stimuli with larger effect sizes have higher test-retest reliability, indicated by the up red arrow, and have the same effect on all sessions, indicated by η. Some active change across sessions could decrease the test-retest reliability, indicated by the down blue arrow. They might have different effects on different sessions, indicated by α_1, α_2, and α_n. The circle represents the measurement error that could decrease the test-retest reliability, indicated by the down blue arrow. There might be different errors on different sessions, indicated by σ_1_, σ_2_, and σ_n_.

However, at the between-person level, the NPS and pain reports’ variances may have been driven by many factors that are irrelevant to the stimuli intensities. One person can report more pain than another because of differences in demographic variables, genetic factors, and psychosocial processes (Fillingim, 2017; Woo & Wager, 2016). For example, individual differences in subjective pain reports might reflect communicative bias, such as “stoics” vs. “communicators.” Meanwhile, the NPS responses might vary due to individual differences in task-related head movement (Engelhardt et al., 2017), respiration (Chang & Glover, 2009; Power et al., 2019), heart rate (Chang et al., 2009), BOLD magnitude (Levin et al., 2001) and inter-individual variation in brain bases for pain reports (Reddan & Wager, 2018). The combination of strong within-person correlations and only modest between-person correlations between the NPS and pain reports indicates that the NPS is not a surrogate for individual differences in pain reports. Instead, the NPS as an objective biological target could be useful for measuring physiological pain in combination with subjective pain reports. For example, a clinical trial is testing a new drug of analgesic. They might want to know how the drug works on pain reports and physiological measures like the NPS. Because pain reports alone may be subject to biases, such as the placebo effects (Tuttle et al., 2015; Zunhammer et al., 2018), the drug may not work in the long term because it does not engage the pain-relevant physiological mechanisms. Laboratory placebo manipulations, on the contrary, only have minimal effects on the NPS (Zunhammer et al., 2018).

Both the NPS (ICC = 0.73 − 0.91) and pain reports (ICC = 0.85 − 0.96) showed excellent short-term (i.e., within one-day) test-retest reliability. Test-retest reliability of pain reports has been extensively examined in previous pain-related studies that showed similar ICC values range from 0.75 − 0.96 (Jackson et al., 2020; Letzen et al., 2014, 2016; Upadhyay et al., 2015). Previous studies have examined the test-retest reliability of univariate brain responses to pain and showed widely varied ICCs in pain-related ROIs (0.32 − 0.88; Letzen et al., 2014; Quiton et al., 2014; Upadhyay et al., 2015), significantly activated clusters (0.33 − 0.74; Jackson et al., 2020) and functional connectivities (−0.17 − 0.77; Letzen et al., 2016).

Compared with the previous univariate brain measures of pain, the NPS showed consistently high performance of short-term test-retest reliability across eight studies. It is noteworthy that although the short-term test-retest reliability is mathematically identical to the internal consistency reliability, they are conceptually different. Internal consistency measures how consistent a set of items, e.g., voxels in NPS, measures a particular construct, e.g., pain (Drost & Others, 2011). At the same time, the short-term test-retest reliability characterizes the short-term temporal stability of measurement, e.g., the NPS response measured within a session (Drost & Others, 2011). High values of internal consistency are not always desirable and could point to the redundancy of items (Streiner, 2003), while high test-retest reliability values are a desirable feature given that the constructs being measured are stable.

To test whether the NPS measure is stable across longer time scales, we examined two studies with 5-day and one-month intervals between sessions. We found that the NPS had high performance in longer-term test-retest reliability when evaluated with sufficient data per person. In our estimation, more than 60 trials per condition are required on average to achieve excellent test-retest reliability, though this is rarely done in practice (Chen et al., 2021; Dang et al., 2020; Rouder & Haaf, 2019). Besides the number of trials per condition, we also tested several other factors influencing the test-retest reliability, including stimulus intensity (e.g., temperature) and whether the measure was calculated in contrast to a baseline or a control condition (see **Figure 2** and **Figure 3(B)**). Previous studies have shown that the test-retest reliabilities of pain reports and activities in ROIs are higher in response to evoked stimuli with higher temperatures (Upadhyay et al., 2015). The test-retest reliabilities of the significantly activated clusters were higher when the measures were calculated in contrast with the baseline than with a control condition (e.g., non-noxious stimuli; Jackson et al., 2020). The current findings of the NPS are consistent with previous studies and are tested quantitatively in a large sample of participants with diverse study procedures. All these factors have a more extensive influence on the NPS than self-reported pain, further supporting the argument that the NPS and pain reports contain different sources of variance (see **Figure 2**).

The complete NPS performance was better than constituent local brain regions for both effect size and test-retest reliability. This finding is consistent with the argument that pain is encoded in distributed brain networks instead of a specific and isolated brain region (Petre et al., 2020; Woo & Wager, 2016). Interestingly, the six regions (i.e., bilateral insula, right dorsal posterior insula, dACC, right S2, and right thalamus) with relatively larger effect sizes and reliabilities were the targets of ascending nociceptive afferents and activated in response to pain stimuli. Other local regions deactivated with pain and are not direct targets of nociceptive afferents have smaller effect sizes and reliabilities (Roy et al., 2012, 2014). The reliabilities of multivariate patterns of ROIs were heterogeneous (i.e., ICCs range from poor to excellent), similar to previous findings of pain-related ROIs using the univariate analyses (Letzen et al., 2014; Quiton et al., 2014; Upadhyay et al., 2015). In contrast, the reliabilities of the complete NPS were more homogeneous and all ranged from good to excellent level across multiple diverse studies.

The current study tests a large number of studies that are diverse in several aspects. Firstly, most of the studies contain some cognitive manipulations along with the painful stimuli, such as cognitive self-regulation intervention to increase or decrease pain (Woo et al., 2015), the combination of painful stimuli with visual or auditory cues for different pain intensities (Atlas et al., 2010; Chang et al., 2015; Jepma et al., 2018; Roy et al., 2014), placebo manipulation (Roy et al., 2014). Secondly, the pain stimuli were applied to different body positions, including arm, foot, leg, and thumbnail, which were supposed to have different sensitivity to pain (Alburquerque-Sendín et al. 2018). Thirdly, the intensities of pain stimuli were largely varied regarding the temperature (44.3 − 50 °C) and duration (1.85 − 12.5 s). The diversities of the studies further support the generalizability of our findings about the measurement properties of the NPS and pain reports. The participants in these studies were mainly young and healthy participants, with only one study testing participants with chronic back pain (i.e., study 10). In study 10, our results showed that the reliability of the NPS in the control group was comparable with other studies when controlling the same number of trials for each participant. The reliabilities of the NPS in the psychotherapy and placebo groups were lower (see **Table S3**). And the reliability of the subjective pain report was lower in study 10 compared with most other studies. We need further research to test the measurement properties of the NPS and subjective pain reports across more diverse participant samples, including clinical populations (Herr et al., 2011; Voepel-Lewis et al., 2010; Walton et al., 2011).

This paper focuses on the NPS because it has been one of the most extensively studied brain signatures for its validity and specificity in the pain domain (Chang et al., 2015; Krishnan et al., 2016; Ma et al., 2016; Van Oudenhove et al., 2020; Wager et al., 2013). The tests in the current study characterizing the reliability, within-person, and between-person variances related to pain reports could be applied to any neuromarker including other pain-related patterns (Brown et al., 2011; Geuter et al., 2020; Kucyi & Davis, 2015; Kutch et al., 2017; López-Solà et al., 2017; Marquand et al., 2010; Woo, Schmidt, et al., 2017). Some other pain signatures possibly could predict individual differences in pain better than the NPS. Our study shows that it is crucial to characterize individual differences across studies and contexts. The correlation with individual differences in pain reports may vary across different experimental instructions and populations. For example, in study 4, we had a selected university population pre-screened for reliable pain reports and pre-calibrated for stimuli intensities and ended up with a very large effect in the between-person level correlation between the NPS and pain reports.

In sum, we find that both the NPS and pain reports have excellent test-retest reliability in a large sample of participants with diverse study procedures. As a measure of individual differences in pain sensitivity, the NPS is only modestly related to pain reports, suggesting that the NPS is not as useful as a surrogate measure of pain report. In contrast, the NPS could serve as an objective biological target to measure physiological pain in combination with subjective pain reports. In the future, other multivariate brain patterns will need to be tested before used as translational biomarkers. Our study provides a blueprint for future studies performing such measurement properties testing and suggests factors that could improve test-retest reliability in future research.

## Notes

### Competing Interest Statement

The authors have declared no competing interest.

## References

Abraham, A., Milham, M. P., Di Martino, A., Craddock, R. C., Samaras, D., Thirion, B., & Varoquaux, G. (2017). Deriving reproducible biomarkers from multi-site resting-state data: An Autism-based example. NeuroImage, 147, 736–745.

Alburquerque-Sendín, F., Madeleine, P., Fernández-de-Las-Peñas, C., Camargo, P. R., & Salvini, T. F. (2018). Spotlight on topographical pressure pain sensitivity maps: A review. Journal of pain research, 11, 215.

Arbabshirani, M. R., Plis, S., Sui, J., & Calhoun, V. D. (2017). Single subject prediction of brain disorders in neuroimaging: Promises and pitfalls. Neuroimage, 145, 137–165.

Ashburner, J., & Friston, K. J. (2005). Unified segmentation. Neuroimage, 26(3), 839–851.

Atlas, L. Y., Bolger, N., Lindquist, M. A., & Wager, T. D. (2010). Brain mediators of predictive cue effects on perceived pain. Journal of Neuroscience, 30(39), 12964–12977.

Atlas, L. Y., Lindquist, M. A., Bolger, N., & Wager, T. D. (2014). Brain mediators of the effects of noxious heat on pain. Pain, 155(8), 1632–1648.

Bakdash, J. Z., & Marusich, L. R. (2017). Repeated Measures Correlation. Frontiers in Psychology, 8, 456.

Barnhart, H. X., Haber, M. J., & Lin, L. I. (2007). An overview on assessing agreement with continuous measurements. Journal of Biopharmaceutical Statistics, 17(4), 529–569.

Bartoshuk, L. M., Duffy, V. B., Green, B. G., Hoffman, H. J., Ko, C. W., Lucchina, L. A., Marks, L. E., Snyder, D. J., & Weiffenbach, J. M. (2004). Valid across-group comparisons with labeled scales: the gLMS versus magnitude matching. Physiology & Behavior, 82(1), 109–114.

Bennett, C. M., & Miller, M. B. (2010). How reliable are the results from functional magnetic resonance imaging? Annals of the New York Academy of Sciences, 1191, 133–155.

Bennett, C. M., & Miller, M. B. (2013). fMRI reliability: influences of task and experimental design. Cognitive, Affective & Behavioral Neuroscience, 13(4), 690–702.

Brown, J. E., Chatterjee, N., Younger, J., & Mackey, S. (2011). Towards a physiology-based measure of pain: patterns of human brain activity distinguish painful from non-painful thermal stimulation. PloS One, 6(9), e24124.

Button, K. S., Ioannidis, J. P. A., Mokrysz, C., Nosek, B. A., Flint, J., Robinson, E. S. J., & Munafò, M. R. (2013). Power failure: why small sample size undermines the reliability of neuroscience. Nature Reviews. Neuroscience, 14(5), 365–376.

Chang, C., Cunningham, J. P., & Glover, G. H. (2009). Influence of heart rate on the BOLD signal: the cardiac response function. NeuroImage, 44(3), 857–869.

Chang, C., & Glover, G. H. (2009). Relationship between respiration, end-tidal CO2, and BOLD signals in resting-state fMRI. NeuroImage, 47(4), 1381–1393.

Chang, L. J., Gianaros, P. J., Manuck, S. B., Krishnan, A., & Wager, T. D. (2015). A Sensitive and Specific Neural Signature for Picture-Induced Negative Affect. PLoS Biology, 13(6), e1002180.

Chen, G., Padmala, S., Chen, Y., Taylor, P. A., Cox, R. W., & Pessoa, L. (2021). To pool or not to pool: Can we ignore cross-trial variability in FMRI?. NeuroImage, 225, 117496.

Cicchetti, D. V., & Sparrow, S. A. (1981). Developing criteria for establishing interrater reliability of specific items: applications to assessment of adaptive behavior. American Journal of Mental Deficiency, 86(2), 127–137.

Cohen, J. (2013). Statistical Power Analysis for the Behavioral Sciences. Academic Press.

Dang, J., King, K. M., & Inzlicht, M. (2020). Why Are Self-Report and Behavioral Measures Weakly Correlated? Trends in Cognitive Sciences, 24(4), 267–269.

Doyle, O. M., Mehta, M. A., & Brammer, M. J. (2015). The role of machine learning in neuroimaging for drug discovery and development. Psychopharmacology, 232(21), 4179–4189.

Drost, E. A., & Others. (2011). Validity and reliability in social science research. Education Research and Perspectives, 38(1), 105.

Dubois, J., & Adolphs, R. (2016). Building a Science of Individual Differences from fMRI. Trends in Cognitive Sciences, 20(6), 425–443.

Elliott, M. L., Knodt, A. R., Cooke, M., Kim, M. J., Melzer, T. R., Keenan, R., Ireland, D., Ramrakha, S., Poulton, R., Caspi, A., Moffitt, T. E., & Hariri, A. R. (2019). General functional connectivity: Shared features of resting-state and task fMRI drive reliable and heritable individual differences in functional brain networks. NeuroImage, 189, 516–532.

Elliott, M. L., Knodt, A. R., Ireland, D., Morris, M. L., Poulton, R., Ramrakha, S., Sison, M. L., Moffitt, T. E., Caspi, A., & Hariri, A. R. (2020). What Is the Test-Retest Reliability of Common Task-Functional MRI Measures? New Empirical Evidence and a Meta-Analysis. Psychological Science, 31(7), 792–806.

Engelhardt, L. E., Roe, M. A., Juranek, J., DeMaster, D., Harden, K. P., Tucker-Drob, E. M., & Church, J. A. (2017). Children’s head motion during fMRI tasks is heritable and stable over time. Developmental Cognitive Neuroscience, 25, 58–68.

Esteban, O., Markiewicz, C. J., Blair, R. W., Moodie, C. A., Isik, A. I., Erramuzpe, A., Kent, J. D., Goncalves, M., DuPre, E., Snyder, M., Oya, H., Ghosh, S. S., Wright, J., Durnez, J., Poldrack, R. A., & Gorgolewski, K. J. (2019). fMRIPrep: a robust preprocessing pipeline for functional MRI. Nature Methods, 16(1), 111–116.

FDA-NIH Biomarker Working Group. (2016). BEST (Biomarkers, EndpointS, and other Tools) Resource. Food and Drug Administration (US).

Fillingim, R. B. (2017). Individual differences in pain: understanding the mosaic that makes pain personal. Pain, 158(Suppl 1), S11.

Gabrieli, J. D. E., Ghosh, S. S., & Whitfield-Gabrieli, S. (2015). Prediction as a humanitarian and pragmatic contribution from human cognitive neuroscience. Neuron, 85(1), 11–26.

Geuter, S., Reynolds Losin, E. A., Roy, M., Atlas, L. Y., Schmidt, L., Krishnan, A., Koban, L., Wager, T. D., & Lindquist, M. A. (2020). Multiple Brain Networks Mediating Stimulus-Pain Relationships in Humans. Cerebral Cortex, 30(7), 4204–4219.

Gordon, E. M., Laumann, T. O., Gilmore, A. W., Newbold, D. J., Greene, D. J., Berg, J. J., Ortega, M., Hoyt-Drazen, C., Gratton, C., Sun, H., Hampton, J. M., Coalson, R. S., Nguyen, A. L., McDermott, K. B., Shimony, J. S., Snyder, A. Z., Schlaggar, B. L., Petersen, S. E., Nelson, S. M., & Dosenbach, N. U. F. (2017). Precision Functional Mapping of Individual Human Brains. Neuron, 95(4), 791–807.e7.

Gratton, C., Kraus, B. T., Greene, D. J., Gordon, E. M., Laumann, T. O., Nelson, S. M., Dosenbach, N. U. F., & Petersen, S. E. (2020). Defining Individual-Specific Functional Neuroanatomy for Precision Psychiatry. Biological Psychiatry, 88(1), 28–39.

Haynes, J. D. (2015). A Primer on Pattern-Based Approaches to fMRI: Principles, Pitfalls, and Perspectives. Neuron, 87(2), 257–270.

Hedge, C., Powell, G., & Sumner, P. (2018). The reliability paradox: Why robust cognitive tasks do not produce reliable individual differences. Behavior Research Methods, 50(3), 1166–1186.

Herr, K., Coyne, P. J., McCaffery, M., Manworren, R., & Merkel, S. (2011). Pain assessment in the patient unable to self-report: position statement with clinical practice recommendations. Pain Management Nursing: Official Journal of the American Society of Pain Management Nurses, 12(4), 230–250.

Herting, M. M., Gautam, P., Chen, Z., Mezher, A., & Vetter, N. C. (2018). Test-retest reliability of longitudinal task-based fMRI: Implications for developmental studies. Developmental Cognitive Neuroscience, 33, 17–26.

Jackson, J. B., O’Daly, O., Makovac, E., Medina, S., Rubio, A. de L., McMahon, S. B., Williams, S. C. R., & Howard, M. A. (2020). Noxious pressure stimulation demonstrates robust, reliable estimates of brain activity and self-reported pain. NeuroImage, 221, 117178.

Jepma, M., Koban, L., van Doorn, J., Jones, M., & Wager, T. D. (2018). Behavioural and neural evidence for self-reinforcing expectancy effects on pain. Nature Human Behaviour, 2(11), 838–855.

Kievit, R. A., Frankenhuis, W. E., Waldorp, L. J., & Borsboom, D. (2013). Simpson’s paradox in psychological science: a practical guide. Frontiers in Psychology, 4, 513.

Koban, L., Jepma, M., López-Solà, M., & Wager, T. D. (2019). Different brain networks mediate the effects of social and conditioned expectations on pain. Nature Communications, 10(1), 4096.

Koo, T. K., & Li, M. Y. (2016). A Guideline of Selecting and Reporting Intraclass Correlation Coefficients for Reliability Research. Journal of Chiropractic Medicine, 15(2), 155–163.

Kraemer, H. C. (2014). The reliability of clinical diagnoses: state of the art. Annual Review of Clinical Psychology, 10, 111–130.

Kragel, P. A., Koban, L., Barrett, L. F., & Wager, T. D. (2018). Representation, Pattern Information, and Brain Signatures: From Neurons to Neuroimaging. Neuron, 99(2), 257–273.

Kragel, P., Han, X., Kraynak, T., Gianaros, P. J., & Wager, T. (2020). fMRI can be highly reliable, but it depends on what you measure. Psyarxiv.

Kriegeskorte, N., Simmons, W. K., Bellgowan, P. S. F., & Baker, C. I. (2009). Circular analysis in systems neuroscience: the dangers of double dipping. Nature Neuroscience, 12(5), 535–540.

Krishnan, A., Woo, C. W., Chang, L. J., Ruzic, L., Gu, X., López-Solà, M., … & Wager, T. D. (2016). Somatic and vicarious pain are represented by dissociable multivariate brain patterns. Elife, 5, e15166.

Kucyi, A., & Davis, K. D. (2015). The dynamic pain connectome. Trends in Neurosciences, 38(2), 86–95.

Kutch, J. J., Ichesco, E., Hampson, J. P., Labus, J. S., Farmer, M. A., Martucci, K. T., Ness, T. J., Deutsch, G., Apkarian, A. V., Mackey, S. C., Klumpp, D. J., Schaeffer, A. J., Rodriguez, L. V., Kreder, K. J., Buchwald, D., Andriole, G. L., Lai, H. H., Mullins, C., Kusek, J. W., … MAPP Research Network. (2017). Brain signature and functional impact of centralized pain: a multidisciplinary approach to the study of chronic pelvic pain (MAPP) network study. Pain, 158(10), 1979–1991.

Letzen, J. E., Boissoneault, J., Sevel, L. S., & Robinson, M. E. (2016). Test-retest reliability of pain-related functional brain connectivity compared with pain self-report. Pain, 157(3), 546–551.

Letzen, J. E., Sevel, L. S., Gay, C. W., O’Shea, A. M., Craggs, J. G., Price, D. D., & Robinson, M. E. (2014). Test-retest reliability of pain-related brain activity in healthy controls undergoing experimental thermal pain. The Journal of Pain: Official Journal of the American Pain Society, 15(10), 1008–1014.

Levin, J. M., Frederick, B. de B., Ross, M. H., Fox, J. F., von Rosenberg, H. L., Kaufman, M. J., Lange, N., Mendelson, J. H., Cohen, B. M., & Renshaw, P. F. (2001). Influence of baseline hematocrit and hemodilution on BOLD fMRI activation. Magnetic Resonance Imaging, 19(8), 1055–1062.

Lindquist, M. A., Krishnan, A., López-Solà, M., Jepma, M., Woo, C. W., Koban, L., … & Wager, T. D. (2017). Group-regularized individual prediction: theory and application to pain. Neuroimage, 145, 274–287.

Lindquist, M. A., Meng Loh, J., Atlas, L. Y., & Wager, T. D. (2009). Modeling the hemodynamic response function in fMRI: efficiency, bias and mis-modeling. NeuroImage, 45(1 Suppl), S187–S198.

López-Solà, M., Woo, C. W., Pujol, J., Deus, J., Harrison, B. J., Monfort, J., & Wager, T. D. (2017). Towards a neurophysiological signature for fibromyalgia. Pain, 158(1), 34–47.

Losin, E. A. R., Woo, C.-W., Medina, N. A., Andrews-Hanna, J. R., Eisenbarth, H., & Wager, T. D. (2020). Author Correction: Neural and sociocultural mediators of ethnic differences in pain. Nature Human Behaviour, 4(6), 656–658.

Manuck, S. B., Brown, S. M., Forbes, E. E., & Hariri, A. R. (2007). Temporal stability of individual differences in amygdala reactivity. The American Journal of Psychiatry, 164(10), 1613–1614.

Marquand, A., Howard, M., Brammer, M., Chu, C., Coen, S., & Mourão-Miranda, J. (2010). Quantitative prediction of subjective pain intensity from whole-brain fMRI data using Gaussian processes. NeuroImage, 49(3), 2178–2189.

Ma, Y., Wang, C., Luo, S., Li, B., Wager, T. D., Zhang, W., Rao, Y., & Han, S. (2016). Serotonin transporter polymorphism alters citalopram effects on human pain responses to physical pain. NeuroImage, 135, 186–196.

McGraw, K. O., & Wong, S. P. (1996). Forming inferences about some intraclass correlation coefficients. Psychological Methods, 1(1), 30–46.

Mumford, J. A., Turner, B. O., Ashby, F. G., & Poldrack, R. A. (2012). Deconvolving BOLD activation in event-related designs for multivoxel pattern classification analyses. NeuroImage, 59(3), 2636–2643.

Nakagawa, S., & Schielzeth, H. (2010). Repeatability for Gaussian and non-Gaussian data: a practical guide for biologists. Biological Reviews, 85(4), 935–956.

Nichols, T. E., Das, S., Eickhoff, S. B., Evans, A. C., Glatard, T., Hanke, M., Kriegeskorte, N., Milham, M. P., Poldrack, R. A., Poline, J.-B., Proal, E., Thirion, B., Van Essen, D. C., White, T., & Yeo, B. T. T. (2017). Best practices in data analysis and sharing in neuroimaging using MRI. Nature Neuroscience, 20(3), 299–303.

Noble, S., Scheinost, D., & Constable, R. T. (2019). A decade of test-retest reliability of functional connectivity: A systematic review and meta-analysis. NeuroImage, 203, 116157.

Nord, C. L., Gray, A., Charpentier, C. J., Robinson, O. J., & Roiser, J. P. (2017). Unreliability of putative fMRI biomarkers during emotional face processing. NeuroImage, 156, 119–127.

O’Connor, D., Potler, N. V., Kovacs, M., Xu, T., Ai, L., Pellman, J., Vanderwal, T., Parra, L. C., Cohen, S., Ghosh, S., Escalera, J., Grant-Villegas, N., Osman, Y., Bui, A., Craddock, R. C., & Milham, M. P. (2017). The Healthy Brain Network Serial Scanning Initiative: a resource for evaluating inter-individual differences and their reliabilities across scan conditions and sessions. GigaScience, 6(2), 1–14.

Orrù, G., Pettersson-Yeo, W., Marquand, A. F., Sartori, G., & Mechelli, A. (2012). Using Support Vector Machine to identify imaging biomarkers of neurological and psychiatric disease: A critical review. Neuroscience and Biobehavioral Reviews, 36(4), 1140–1152.

Pannunzi, M., Hindriks, R., Bettinardi, R. G., Wenger, E., Lisofsky, N., Martensson, J., Butler, O., Filevich, E., Becker, M., Lochstet, M., Kühn, S., & Deco, G. (2017). Resting-state fMRI correlations: From link-wise unreliability to whole brain stability. NeuroImage, 157, 250–262.

Petre, B., Kragel, P. A., Atlas, L. Y., Geuter, S., Jepma, M., Koban, L., … & Wager, T. D. (2020). Evoked pain intensity representation is distributed across brain systems: A multistudy mega-analysis. BioRxiv.

Plichta, M. M., Schwarz, A. J., Grimm, O., Morgen, K., Mier, D., Haddad, L., Gerdes, A. B. M., Sauer, C., Tost, H., Esslinger, C., Colman, P., Wilson, F., Kirsch, P., & Meyer-Lindenberg, A. (2012). Test–retest reliability of evoked BOLD signals from a cognitive–emotive fMRI test battery. NeuroImage, 60(3), 1746–1758.

Poldrack, R. A., Baker, C. I., Durnez, J., Gorgolewski, K. J., Matthews, P. M., Munafò, M. R., … & Yarkoni, T. (2017). Scanning the horizon: towards transparent and reproducible neuroimaging research. Nature reviews neuroscience, 18(2), 115.

Power, J. D., Lynch, C. J., Silver, B. M., Dubin, M. J., Martin, A., & Jones, R. M. (2019). Distinctions among real and apparent respiratory motions in human fMRI data. NeuroImage, 201, 116041.

Quiton, R. L., Keaser, M. L., Zhuo, J., Gullapalli, R. P., & Greenspan, J. D. (2014). Intersession reliability of fMRI activation for heat pain and motor tasks. NeuroImage. Clinical, 5, 309–321.

Reddan, M. C., Lindquist, M. A., & Wager, T. D. (2017). Effect Size Estimation in Neuroimaging. JAMA Psychiatry, 74(3), 207–208.

Reddan, M. C., & Wager, T. D. (2018). Modeling Pain Using fMRI: From Regions to Biomarkers. Neuroscience Bulletin, 34(1), 208–215.

Rissman, J., Gazzaley, A., & D’Esposito, M. (2004). Measuring functional connectivity during distinct stages of a cognitive task. NeuroImage, 23(2), 752–763.

Rouder, J. N., & Haaf, J. M. (2019). A psychometrics of individual differences in experimental tasks. Psychonomic Bulletin & Review, 26(2), 452–467.

Roy, M., Shohamy, D., Daw, N., Jepma, M., Wimmer, G. E., & Wager, T. D. (2014). Representation of aversive prediction errors in the human periaqueductal gray. Nature Neuroscience, 17(11), 1607–1612.

Roy, M., Shohamy, D., & Wager, T. D. (2012). Ventromedial prefrontal-subcortical systems and the generation of affective meaning. Trends in Cognitive Sciences, 16(3), 147–156.

Sawilowsky, S. S. (2009). New effect size rules of thumb. Journal of Modern Applied Statistical Methods, 8(2), 26.

Shmuel, A., Chaimow, D., Raddatz, G., Ugurbil, K., & Yacoub, E. (2010). Mechanisms underlying decoding at 7 T: ocular dominance columns, broad structures, and macroscopic blood vessels in V1 convey information on the stimulated eye. Neuroimage, 49(3), 1957–1964.

Shrout, P. E., & Fleiss, J. L. (1979). Intraclass correlations: uses in assessing rater reliability. Psychological Bulletin, 86(2), 420–428.

Streiner, D. L. (2003). Starting at the beginning: an introduction to coefficient alpha and internal consistency. Journal of Personality Assessment, 80(1), 99–103.

Tuttle, A. H., Tohyama, S., Ramsay, T., Kimmelman, J., Schweinhardt, P., Bennett, G. J., & Mogil, J. S. (2015). Increasing placebo responses over time in U.S. clinical trials of neuropathic pain. Pain, 156(12), 2616–2626.

Upadhyay, J., Lemme, J., Anderson, J., Bleakman, D., Large, T., Evelhoch, J. L., Hargreaves, R., Borsook, D., & Becerra, L. (2015). Test-retest reliability of evoked heat stimulation BOLD fMRI. Journal of Neuroscience Methods, 253, 38–46.

Van Oudenhove, L., Kragel, P. A., Dupont, P., Ly, H. G., Pazmany, E., Enzlin, P., Rubio, A., Delon-Martin, C., Bonaz, B., Aziz, Q., Tack, J., Fukudo, S., Kano, M., & Wager, T. D. (2020). Common and distinct neural representations of aversive somatic and visceral stimulation in healthy individuals. Nature Communications, 11(1), 5939.

Voepel-Lewis, T., Zanotti, J., Dammeyer, J. A., & Merkel, S. (2010). Reliability and validity of the face, legs, activity, cry, consolability behavioral tool in assessing acute pain in critically ill patients. American journal of critical care, 19(1), 55–61.

Wager, T. D., Atlas, L. Y., Lindquist, M. A., Roy, M., Woo, C. W., & Kross, E. (2013). An fMRI-based neurologic signature of physical pain. The New England Journal of Medicine, 368(15), 1388–1397.

Wager, T. D., & Nichols, T. E. (2003). Optimization of experimental design in fMRI: a general framework using a genetic algorithm. Neuroimage, 18(2), 293–309.

Walton, D. M., Macdermid, J. C., Nielson, W., Teasell, R. W., Chiasson, M., & Brown, L. (2011). Reliability, standard error, and minimum detectable change of clinical pressure pain threshold testing in people with and without acute neck pain. The Journal of Orthopaedic and Sports Physical Therapy, 41(9), 644–650.

Woo, C. W., Chang, L. J., Lindquist, M. A., & Wager, T. D. (2017). Building better biomarkers: brain models in translational neuroimaging. Nature Neuroscience, 20(3), 365–377.

Woo, C. W., Roy, M., Buhle, J. T., & Wager, T. D. (2015). Distinct brain systems mediate the effects of nociceptive input and self-regulation on pain. PLoS Biology, 13(1), e1002036.

Woo, C.-W., Schmidt, L., Krishnan, A., Jepma, M., Roy, M., Lindquist, M. A., Atlas, L. Y., & Wager, T. D. (2017). Quantifying cerebral contributions to pain beyond nociception. Nature Communications, 8, 14211.

Woo, C. W., & Wager, T. D. (2016). What reliability can and cannot tell us about pain report and pain neuroimaging. Pain, 157(3), 511–513.

Assessing Variations in Areal Organization for the Intrinsic Brain: From Fingerprints to Reliability. Cerebral Cortex, 26(11), 4192–4211.

Yoo, K., Rosenberg, M. D., Noble, S., Scheinost, D., Constable, R. T., & Chun, M. M. (2019). Multivariate approaches improve the reliability and validity of functional connectivity and prediction of individual behaviors. NeuroImage, 197, 212–223.

Zunhammer, M., Bingel, U., Wager, T. D., & Placebo Imaging Consortium. (2018). Placebo Effects on the Neurologic Pain Signature: A Meta-analysis of Individual Participant Functional Magnetic Resonance Imaging Data. JAMA Neurology, 75(11), 1321–1330.

Zuo, X.-N., & Xing, X.-X. (2014). Test-retest reliabilities of resting-state FMRI measurements in human brain functional connectomics: a systems neuroscience perspective. Neuroscience and Biobehavioral Reviews, 45, 100–118.

Zuo, X.-N., Xu, T., & Milham, M. P. (2019). Harnessing reliability for neuroscience research. Nature Human Behaviour, 3(8), 768–771.

