## Supplementary information for "Effect sizes and test-retest reliability of the fMRI-based Neurologic Pain Signature"

**Table S1.** Inferential statistics for four types of NPS effects in Studies 1-8

| **Study** | **Mean response of NPS** | | | | **Within-subject correlation with temperature** | | | | | **Within-subject correlation with pain** | | | | | **Between-subject correlation with pain** | | |
|---|---|---|---|---|---|---|---|---|---|---|---|---|---|---|---|---|---|
| | **t** | **df** | **p** | **d** | **within-r** | **t** | **df** | **p** | **d** | **within-r** | **t** | **df** | **p** | **d** | **between-r** | **p** | **d** |
| Study1 | 10.19 | 32 | <0.001 | 1.77 | 0.27 | 6.25 | 32 | <0.001 | 1.09 | 0.28 | 6.39 | 32 | <0.001 | 1.11 | -0.13 | 0.46 | -0.27 |
| Study2 | 13.86 | 27 | <0.001 | 2.62 | 0.30 | 10.98 | 27 | <0.001 | 2.07 | 0.33 | 8.69 | 27 | <0.001 | 1.64 | 0.16 | 0.40 | 0.33 |
| Study3 | 19.22 | 92 | <0.001 | 1.99 | 0.22 | 9.27 | 92 | <0.001 | 0.96 | 0.23 | 9.83 | 91 | <0.001 | 1.02 | 0.04 | 0.70 | 0.08 |
| Study4 | 5.02 | 16 | <0.001 | 1.22 | 0.31 | 6.91 | 16 | <0.001 | 1.68 | 0.34 | 6.58 | 16 | <0.001 | 1.59 | 0.74 | <0.001 | 2.20 |
| Study5 | 17.07 | 49 | <0.001 | 2.41 | 0.42 | 18.91 | 49 | <0.001 | 2.67 | 0.35 | 13.56 | 49 | <0.001 | 1.92 | 0.19 | 0.18 | 0.39 |
| Study6 | 6.97 | 18 | <0.001 | 1.60 | 0.05 | 2.32 | 18 | 0.03 | 0.53 | 0.14 | 5.22 | 18 | <0.001 | 1.20 | 0.31 | 0.20 | 0.65 |
| Study7 | 8.91 | 28 | <0.001 | 1.65 | 0.30 | 11.65 | 28 | <0.001 | 2.16 | 0.35 | 11.49 | 28 | <0.001 | 2.13 | 0.04 | 0.86 | 0.07 |
| Study8 | 10.60 | 25 | <0.001 | 2.08 | 0.13 | 4.20 | 25 | <0.001 | 0.82 | 0.17 | 4.81 | 25 | <0.001 | 0.94 | 0.22 | 0.27 | 0.46 |
| **Mean** | | | | **1.92** | | | | | **1.50** | | | | | **1.45** | | | **0.49** |

**Note**: One-sample t-tests treat the subject as the unit of observation (i.e., the subject is a random effect). In between-subject correlation, transformation between r and cohen's d, i.e., d = $\frac{2\times r}{\sqrt{1-r^2}}$.

**Table S2**. Inferential statistics comparing effect size of whole-NPS with local NPS responses in Studies 1-8

| | Rank | Mean response | | | | Within-subject correlation with temperature | | | | Within-subject correlation with pain | | | | Between-subject correlation with pain | | | |
|---|---|---|---|---|---|---|---|---|---|---|---|---|---|---|---|---|---|
| | | **Brain region** | **Mean (se)** | **t** | **q-fdr** | **Brain region** | **Mean (se)** | **t** | **q-fdr** | **Brain region** | **Mean (se)** | **t** | **q-fdr** | **Brain region** | **Mean (se)** | **t** | **q-fdr** |
| | | **NPS** | 1.92 (0.16) | | | **NPS** | 1.50 (0.27) | | | **NPS** | 1.45 (0.16) | | | **NPS** | 0.49 (0.26) | | |
| **Positive regions** | **Rank 1** | **rmIns** | 1.72 (0.19) | 1.63 | 0.148 | **rmIns** | 1.21 (0.26) | 1.92 | 0.097 | **dACC** | 1.19 (0.12) | 2.19 | 0.065 | **rdpIns** | 0.40 (0.31) | 0.65 | 0.539 |
| | **Rank 2** | **rdpIns** | 1.43 (0.11) | 3.79 | 0.007 | **dACC** | 1.12 (0.22) | 2.65 | 0.033 | **rmIns** | 1.14 (0.14) | 3.98 | 0.005 | **rmIns** | 0.37 (0.14) | 0.67 | 0.526 |
| | **Rank 3** | **rS2** | 1.21 (0.15) | 5.94 | 0.001 | **rdpIns** | 1.00 (0.16) | 3.56 | 0.009 | **lmIns** | 1.04 (0.10) | 4.52 | 0.003 | **vermis** | 0.32 (0.11) | 0.77 | 0.469 |
| | **Rank 4** | **lmIns** | 1.19 (0.18) | 6.18 | 0.001 | **lmIns** | 0.92 (0.18) | 4.36 | 0.003 | **rdpIns** | 1.02 (0.14) | 4.96 | 0.002 | **rThal** | 0.30 (0.16) | 1.11 | 0.304 |
| | **Rank 5** | **dACC** | 1.01 (0.18) | 9.26 | <0.001 | **rS2** | 0.76 (0.15) | 3.68 | 0.008 | **rThal** | 0.85 (0.08) | 6.33 | <0.001 | **lmIns** | 0.27 (0.11) | 0.99 | 0.354 |
| | **Rank 6** | **rThal** | 0.70 (0.16) | 6.99 | <0.001 | **rThal** | 0.74 (0.14) | 4.22 | 0.004 | **rS2** | 0.83 (0.09) | 4.94 | 0.002 | **dACC** | 0.26 (0.18) | 1.21 | 0.266 |
| | **Rank 7** | **vermis** | 0.44 (0.11) | 10.04 | <0.001 | **vermis** | 0.63 (0.15) | 3.98 | 0.005 | **vermis** | 0.80 (0.11) | 4.03 | 0.005 | **rS2** | 0.24 (0.20) | 1.98 | 0.088 |
| | **Rank 8** | **rV1** | -0.61 (0.28) | 8.02 | <0.001 | **rV1** | 0.17 (0.20) | 3.97 | 0.005 | **rV1** | 0.19 (0.20) | 4.24 | 0.004 | **rV1** | 0.21 (0.08) | 1.21 | 0.265 |
| **Negative regions** | **Rank 1** | **rIPL** | 0.73 (0.24) | 4.42 | 0.004 | **rIPL** | 0.24 (0.20) | 3.95 | 0.006 | **rIPL** | 0.17 (0.21) | 6.42 | <0.001 | **pgACC** | 0.10 (0.15) | 1.64 | 0.144 |
| | **Rank 2** | **pgACC** | 0.60 (0.19) | 5.59 | 0.002 | **lSTS** | 0.14 (0.20) | 4.53 | 0.003 | **pgACC** | 0.15 (0.14) | 7.03 | <0.001 | **rLOC** | 0.08 (0.15) | 2.11 | 0.072 |
| | **Rank 3** | **PCC** | 0.51 (0.28) | 3.86 | 0.007 | **pgACC** | 0.09 (0.15) | 4.44 | 0.003 | **lSTS** | 0.05 (0.20) | 7.07 | <0.001 | **rpLOC** | -0.01 (0.12) | 1.46 | 0.187 |
| | **Rank 4** | **lLOC** | 0.44 (0.27) | 4.80 | 0.003 | **rLOC** | 0.06 (0.19) | 4.69 | 0.002 | **rpLOC** | 0.03 (0.13) | 9.81 | <0.001 | **lLOC** | -0.01 (0.15) | 2.08 | 0.076 |
| | **Rank 5** | **rpLOC** | 0.34 (0.26) | 5.41 | 0.002 | **rpLOC** | 0.04 (0.14) | 4.81 | 0.002 | **lOLC** | 0.02 (0.16) | 8.50 | <0.001 | **lSTS** | -0.06 (0.11) | 1.84 | 0.108 |
| | **Rank 6** | **rLOC** | 0.33 (0.26) | 5.23 | 0.002 | **lLOC** | 0.02 (0.19) | 4.53 | 0.003 | **rLOC** | -0.01 (0.21) | 7.56 | <0.001 | **PCC** | -0.26 (0.17) | 2.12 | 0.072 |
| | **Rank 7** | **lSTS** | 0.31 (0.11) | 8.71 | <0.001 | **PCC** | -0.21 (0.24) | 4.54 | 0.003 | **PCC** | -0.33 (0.19) | 9.66 | <0.001 | **rIPL** | -0.27 (0.17) | 2.83 | 0.025 |

**Note**: Paired t-tests between effect size of NPS and other brain regions treat study as the unit of observation (i.e., study is a random effect). To adjust the false positive rate in multiple comparisons, FDR corrected q values are reported. Ins denotes Insula, V1 primary visual area, S2 secondary somatosensory cortex, ACC anterior cingulate cortex, Thal thalamus, STS superior temporal sulcus, PCC posterior cingulate cortex, LOC lateral occipital complex, and IPL inferior parietal lobule. Direction is indicated with preceding lowercase letters as follows: r denotes right, l left, m middle, d dorsal, p posterior, pg perigenual.

**Table S3**. Short-term and long-term test-retest reliability of NPS and pain reports

| Study | Type | NPS | | Pain reports | |
|---|---|---|---|---|---|
| | | **ICC** | **95CI** | **ICC** | **95CI** |
| Study1 | ICC(3,k) | 0.87 | 0.77 - 0.93 | 0.95 | 0.91 - 0.97 |
| Study2 | ICC(3,k) | 0.88 | 0.79 - 0.94 | 0.89 | 0.80 - 0.95 |
| Study3 | ICC(3,k) | 0.91 | 0.88 - 0.94 | 0.92 | 0.89 - 0.94 |
| Study4 | ICC(3,k) | 0.73 | 0.38 - 0.89 | 0.85 | 0.65 - 0.94 |
| Study5 | ICC(3,k) | 0.87 | 0.80 - 0.92 | 0.96 | 0.93 - 0.97 |
| Study6 | ICC(3,k) | 0.85 | 0.66 - 0.93 | 0.93 | 0.85 - 0.97 |
| Study7 | ICC(3,k) | 0.87 | 0.76 - 0.93 | 0.93 | 0.88 - 0.96 |
| Study8 | ICC(3,k) | 0.75 | 0.52 - 0.87 | 0.92 | 0.84 - 0.96 |
| Study9 | ICC(3,1) | 0.74 | 0.61 - 0.84 | 0.87 | 0.80 - 0.92 |
| *Study10 - control group | ICC(3,1) | 0.46 | 0.22 - 0.65 | 0.26 | -0.15 - 0.49 |
| *Study10 - placebo group | ICC(3,1) | 0.17 | -0.01 - 0.41 | 0.27 | 0.02 - 0.50 |
| *Study10 - psychotherapy group | ICC(3,1) | 0.34 | 0.09 - 0.55 | 0.18 | -0.08 - 0.41 |

* In Study 10, we reported the control group results in the main text.

**Table S4**. Inferential statistics comparing the reliability of whole-NPS with local NPS responses in Studies 1-8

| | Rank | Brain | Mean (se) | t | q-fdr |
|---|---|---|---|---|---|
| | | **NPS** | 0.84 (0.02) | | |
| **Positive Region** | **Rank 1** | rS2 | 0.83 (0.03) | 0.70 | 0.508 |
| | **Rank 2** | rmIns | 0.80 (0.04) | 1.48 | 0.182 |
| | **Rank 3** | rV1 | 0.78 (0.05) | 1.32 | 0.229 |
| | **Rank 4** | rdpIns | 0.75 (0.06) | 2.12 | 0.072 |
| | **Rank 5** | dACC | 0.75 (0.08) | 1.57 | 0.161 |
| | **Rank 6** | lmIns | 0.72 (0.11) | 1.26 | 0.247 |
| | **Rank 7** | rThal | 0.61 (0.08) | 3.72 | 0.007 |
| | **Rank 8** | vermis | 0.58 (0.16) | 1.84 | 0.109 |
| **Negative Region** | **Rank 1** | lSTS | 0.72 (0.07) | 1.99 | 0.086 |
| | **Rank 2** | PCC | 0.70 (0.08) | 2.31 | 0.054 |
| | **Rank 3** | lLOC | 0.64 (0.11) | 2.24 | 0.060 |
| | **Rank 4** | rpLOC | 0.63 (0.14) | 1.63 | 0.147 |
| | **Rank 5** | rIPL | 0.63 (0.10) | 2.37 | 0.050 |
| | **Rank 6** | rLOC | 0.60 (0.16) | 1.75 | 0.123 |
| | **Rank 7** | pgACC | 0.58 (0.08) | 4.27 | 0.004 |

**Note**: Paired t-tests of reliability, i.e., ICC(3,k), between NPS and other brain regions treat study as the unit of observation (i.e., study is a random effect). To adjust the false positive rate in multiple comparisons, FDR corrected q values are reported. Ins denotes Insula, V1 primary visual area, S2 secondary somatosensory cortex, ACC anterior cingulate cortex, Thal thalamus, STS superior temporal sulcus, PCC posterior cingulate cortex, LOC lateral occipital complex, and IPL inferior parietal lobule. Direction is indicated with preceding lowercase letters as follows: r denotes right, l left, m middle, d dorsal, p posterior, pg perigenual.

**Table S5**. Comparison of NPS performance in effect size and reliability among three computation methods in Studies 1-8

| Index | Mean response | | | Within-subject correlation with temperature | | | Within-subject correlation with pain | | | Between-subject correlation with pain | | | Reliability | | |
|---|---|---|---|---|---|---|---|---|---|---|---|---|---|---|---|
| | **Mean (se)** | **F** | **p** | **Mean (se)** | **F** | **p** | **Mean (se)** | **F** | **p** | **Mean (se)** | **F** | **p** | **Mean (se)** | **F** | **p** |
| **Dot product** | 1.92 (0.16) | 0.87 | 0.43 | 1.50 (0.27) | 0.30 | 0.74 | 1.45 (0.16) | 0.70 | 0.51 | 0.49 (0.26) | 0.18 | 0.83 | 0.84 (0.02) | 0.03 | 0.98 |
| **correlation** | 2.28 (0.23) | | | 1.27 (0.22) | | | 1.17 (0.20) | | | 0.29 (0.24) | | | 0.84 (0.02) | | |
| **cosine** | 2.19 (0.21) | | | 1.27 (0.23) | | | 1.19 (0.19) | | | 0.31 (0.26) | | | 0.84 (0.02) | | |

**Note**: One-way ANOVAs of NPS effect sizes and reliability, i.e., ICC(3,k), treat the study as the unit of observation (i.e., the study is a random effect).

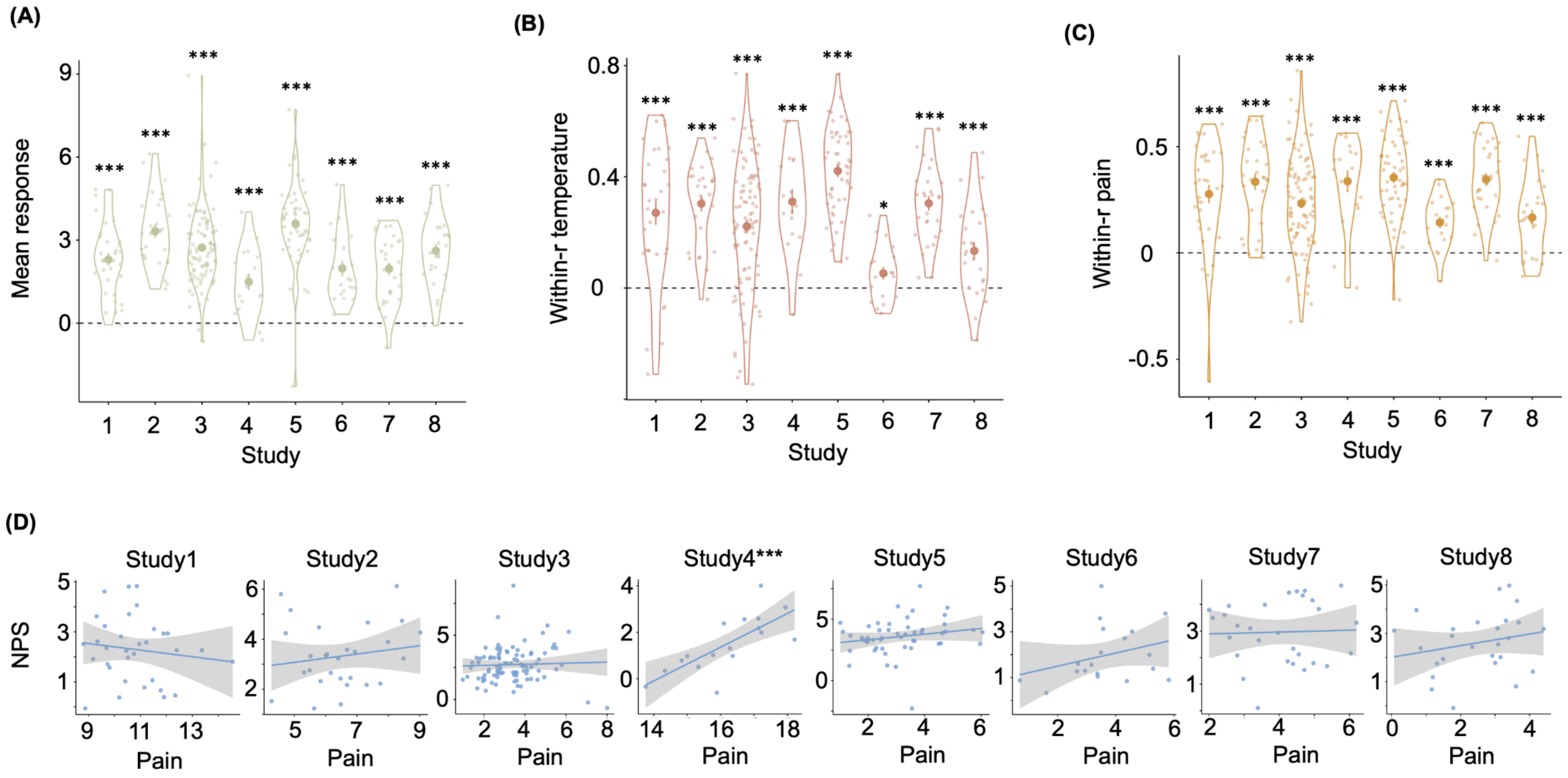


**Figure S1. Four tests of NPS in response to pain stimuli**. **(A) Mean response of NPS.** Each big dot represents the mean response of NPS in each study; the vertical bar represents the standard error; each small dot represents the mean NPS response of one participant, and the violin plot represents the distribution of all participants in each study. For display only and make the NPS values comparable across studies, we rescaled the NPS response by dividing the mean of the absolute deviations from the mean of all studies (using the *mad* function in MATLAB). **(B) Within-person correlation between the NPS response and the temperature.** Each big dot represents the mean r-value of the correlation between the NPS response and the temperature; the vertical bar represents the standard error; each small dot represents the r-value of one participant, and the violin plot represents the distribution of all participants in each study. **(C) Within-person correlation between the NPS response and the subjective pain reports.** Each big dot represents the mean r-value of the correlation between the NPS response and the subjective pain reports; the vertical bar represents the standard error; each small dot represents the r-value of one participant, and the violin plot represents the distribution of all participants in each study. **(D) Between-person correlation between the NPS response and participants' mean subjective pain reports.** Each dot represents one participant; the line represents the linear relationship between the mean of the subjective pain reports and the mean of the NPS response of each participant, and the shadow represents the standard error. The NPS response and pain ratings were rescaled by dividing the mean of the absolute deviations from the mean of all studies (using the *mad* function in MATLAB). *** $p < 0.001$; ** $p<0.005$; * $p<0.05$.

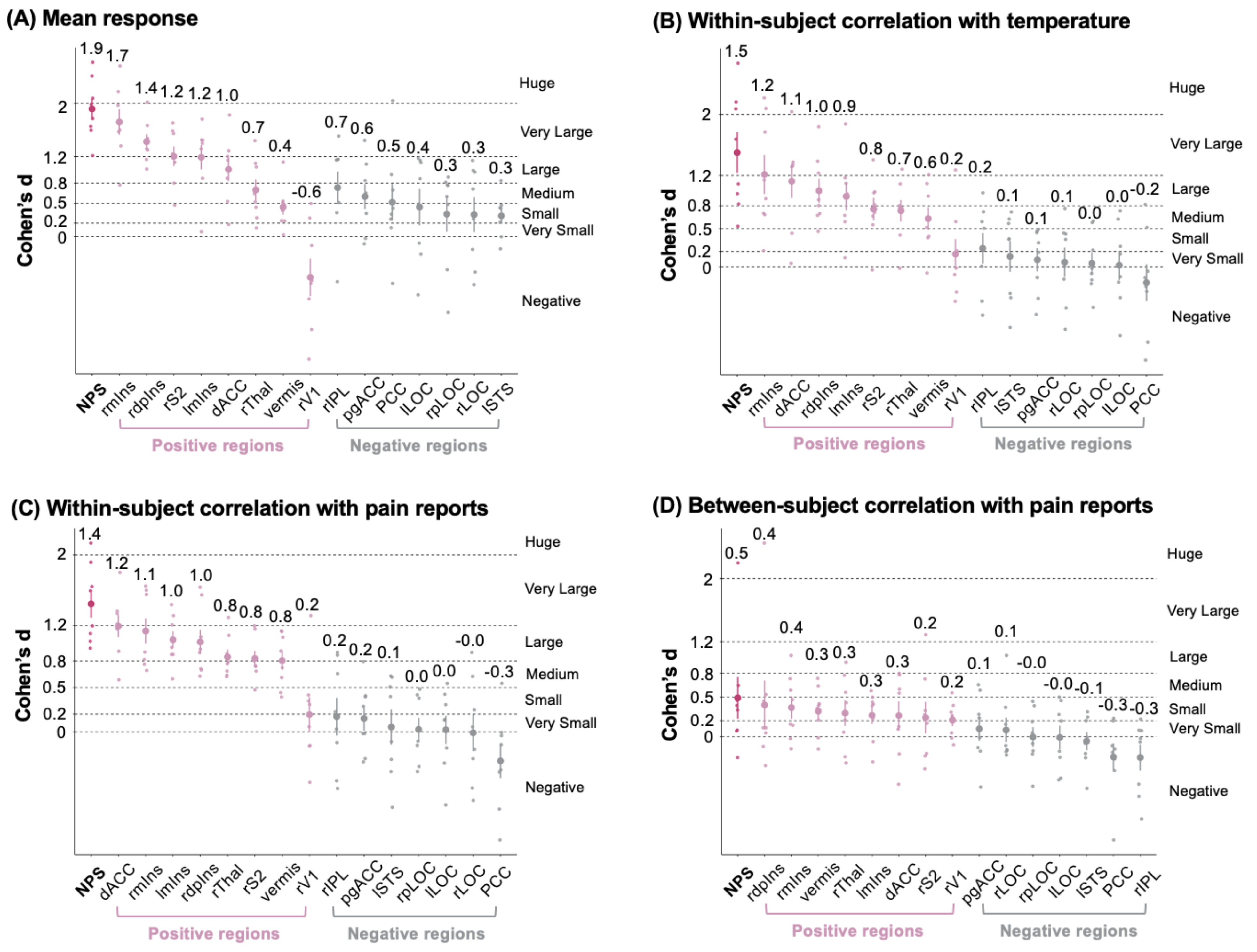


**Figure S2. Four types of the effect size of NPS local brain regions.** (A) The effect size of the mean response of NPS and local areas. (B) The effect size of the within-person correlation between brain and the temperature. (C) The effect size of the within-person correlation between brain and the subjective pain ratings. (D) The effect size of the between-subject correlation between brain and the subjective pain ratings. Each big dot represents the mean effect size across studies. Each small dot represents the effect size for an individual study (Studies 1 to 8). The vertical bar represents the standard error. Ins denotes Insula, V1 primary visual area, S2 secondary somatosensory cortex, ACC anterior cingulate cortex, Thal thalamus, STS superior temporal sulcus, PCC posterior cingulate cortex, LOC lateral occipital complex, and IPL inferior parietal lobule. Direction is indicated with preceding lowercase letters as follows: r denotes right, l left, m middle, d dorsal, p posterior, pg perigenual.

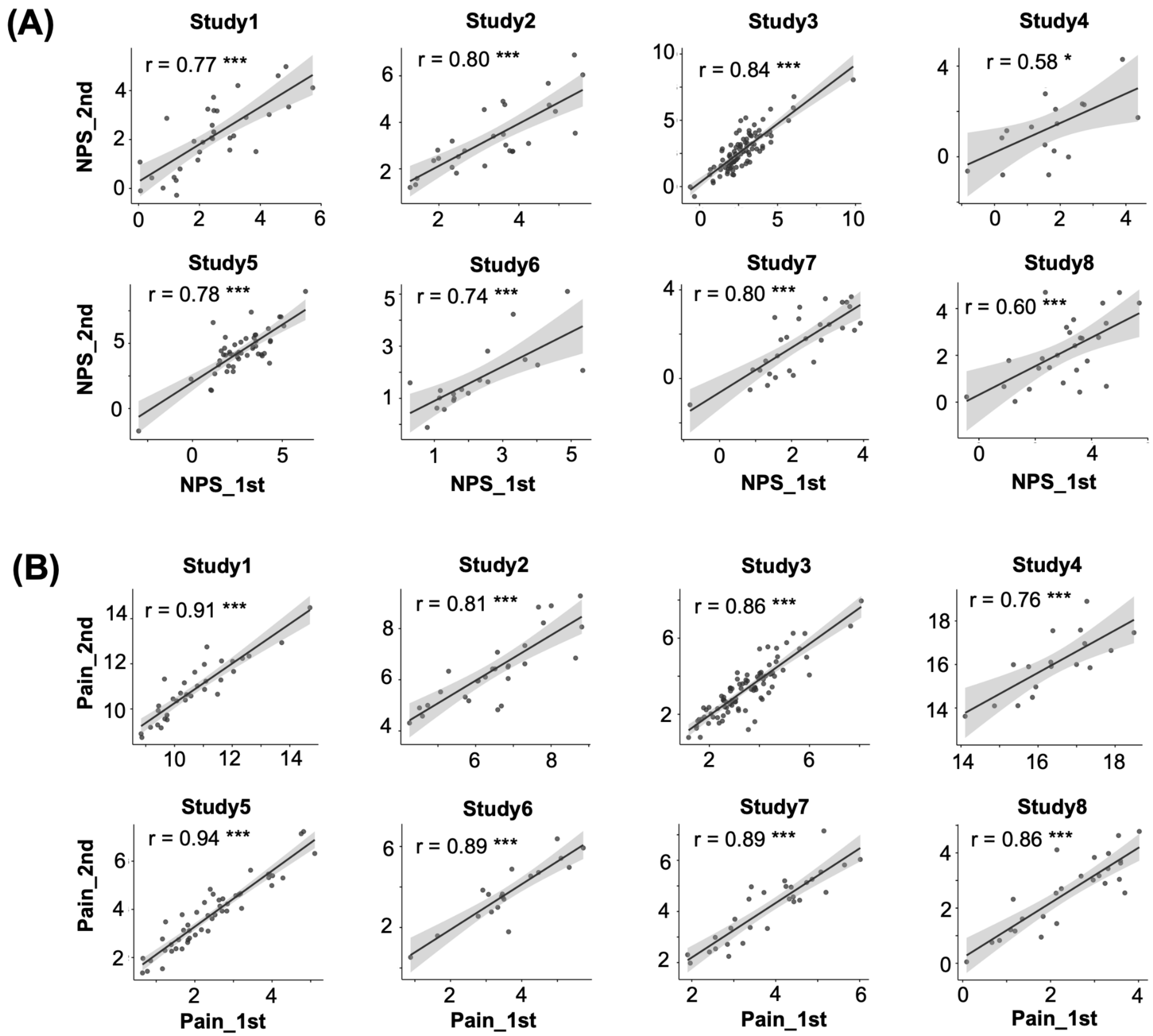


**Figure S3. Illustration of short-term test-retest reliability of the NPS and subjective pain reports.** (A) Correlation between the averaged NPS response of the first-half and second-half trials. (B) Correlation between the averaged subjective pain reports of the first-half and second-half trials. Each dot represents one participant; the line represents the linear relationship between the averaged measurements of the first-half and second-half trials. The NPS response and pain ratings were rescaled by dividing the mean of the absolute deviations from the mean of all studies (using the *mad* function in MATLAB). *** p < 0.001; ** p<0.005; * p<0.05.
